# KBTBD4 Cancer Hotspot Mutations Drive Neomorphic Degradation of HDAC1/2 Corepressor Complexes

**DOI:** 10.1101/2024.05.14.593970

**Authors:** Xiaowen Xie, Olivia Zhang, Megan J.R. Yeo, Ceejay Lee, Stefan A. Harry, Leena Paul, Yiran Li, N. Connor Payne, Eunju Nam, Hui Si Kwok, Hanjie Jiang, Haibin Mao, Jennifer L. Hadley, Hong Lin, Melissa Batts, Pallavi M. Gosavi, Vincenzo D’Angiolella, Philip A. Cole, Ralph Mazitschek, Paul A. Northcott, Ning Zheng, Brian B. Liau

## Abstract

Cancer mutations can create neomorphic protein-protein interactions to drive aberrant function^1^. As a substrate receptor of the CULLIN3-RBX1 E3 ubiquitin ligase complex, KBTBD4 is recurrently mutated in medulloblastoma (MB)^2^, the most common embryonal brain tumor in children, and pineoblastoma^3^. These mutations impart gain-of-function to KBTBD4 to induce aberrant degradation of the transcriptional corepressor CoREST^4^. However, their mechanism of action remains unresolved. Here, we elucidate the mechanistic basis by which KBTBD4 mutations promote CoREST degradation through engaging HDAC1/2, the direct neomorphic target of the substrate receptor. Using deep mutational scanning, we systematically map the mutational landscape of the KBTBD4 cancer hotspot, revealing distinct preferences by which insertions and substitutions can promote gain-of-function and the critical residues involved in the hotspot interaction. Cryo-electron microscopy (cryo-EM) analysis of two distinct KBTBD4 cancer mutants bound to LSD1-HDAC1-CoREST reveals that a KBTBD4 homodimer asymmetrically engages HDAC1 with two KELCH-repeat propeller domains. The interface between HDAC1 and one of the KBTBD4 propellers is stabilized by the MB mutations, which directly insert a bulky side chain into the active site pocket of HDAC1. Our structural and mutational analyses inform how this hotspot E3-neo-substrate interface can be chemically modulated. First, our results unveil a converging shape complementarity-based mechanism between gain-of-function E3 mutations and a molecular glue degrader, UM171. Second, we demonstrate that HDAC1/2 inhibitors can block the mutant KBTBD4-HDAC1 interface, the aberrant degradation of CoREST, and the growth of KBTBD4-mutant MB models. Altogether, our work reveals the structural and mechanistic basis of cancer mutation-driven neomorphic protein-protein interactions and pharmacological strategies to modulate their action for therapeutic applications.

## Introduction

Human genetic variation and somatic mutations in protein-coding genes can alter their protein-protein interactions (PPIs) to drive disease states^1,5,6^. While many of these mutations cause loss-of-function, recent studies have demonstrated how they can also promote neomorphic PPIs with aberrant functions^6–9^. Understanding the molecular mechanisms governing how mutations can enable ‘neo-PPIs’ will be critical for not only understanding disease etiology but also for guiding therapeutic approaches, such as molecular glues and PPI inhibitors, to chemically modulate these interfaces^10,11^.

In the ubiquitin-proteasome system, human disease mutations have long been known to compromise the functions of several E3 ubiquitin ligases^12–15^. By contrast, gain-of-function mutations in E3s promoting aberrant degradation of substrate proteins have only recently emerged as a fascinating phenomenon. While cases of hypermorphic E3 mutations are better documented, leading to unscheduled substrate ubiquitination by altering ligase stability or regulation^16–19^, neomorphic E3 mutations that directly induce neo-substrate engagement and degradation represent a new paradigm in E3 ligase dysregulation.

Cancer mutations in *KBTBD4*, a CULLIN3-RBX1 E3 ligase (CRL3) substrate receptor, present the first compelling case of E3 ligase neomorphic mutations. *KBTBD4* is recurrently mutated in Group 3 and 4 MBs^2^, molecular subtypes associated with poor outcomes and lacking effective treatment options. These mutations occur in a hotspot in the 2b-2c loop of the KELCH-repeat β-propeller and comprise remarkable molecular diversity, spanning 1-5 amino acid insertion-deletions (indel or delins) as well as point substitutions^2^ (**Fig. 1a**). The most common mutations include P311delinsPP and R313delinsPRR (abbreviated as P and PR, hereafter), which can promote the neomorphic degradation of the CoREST and LSD1 subunits of the LSD1-HDAC1/2-CoREST (LHC) complex^4^. Remarkably, KBTBD4 has also been identified as the target of UM171, a small molecule agonist of hematopoietic stem cell expansion that induces CoREST degradation^20,21^. Nevertheless, the exact molecular target and mechanism of mutant KBTBD4 remains unclear. Here we elucidate the structural basis and mechanism of KBTBD4 MB mutations, resolving how the mutations promote neomorphic E3 ligase complexation to the LHC complex via HDAC1/2 to induce selective corepressor degradation. Furthermore, our results establish a striking, converging mechanism between cancer mutations and a molecular glue degrader and uncover previously unrealized therapeutic opportunities for treating *KBTBD4*-mutant MBs with HDAC1/2 inhibitors.

**Figure 1.**
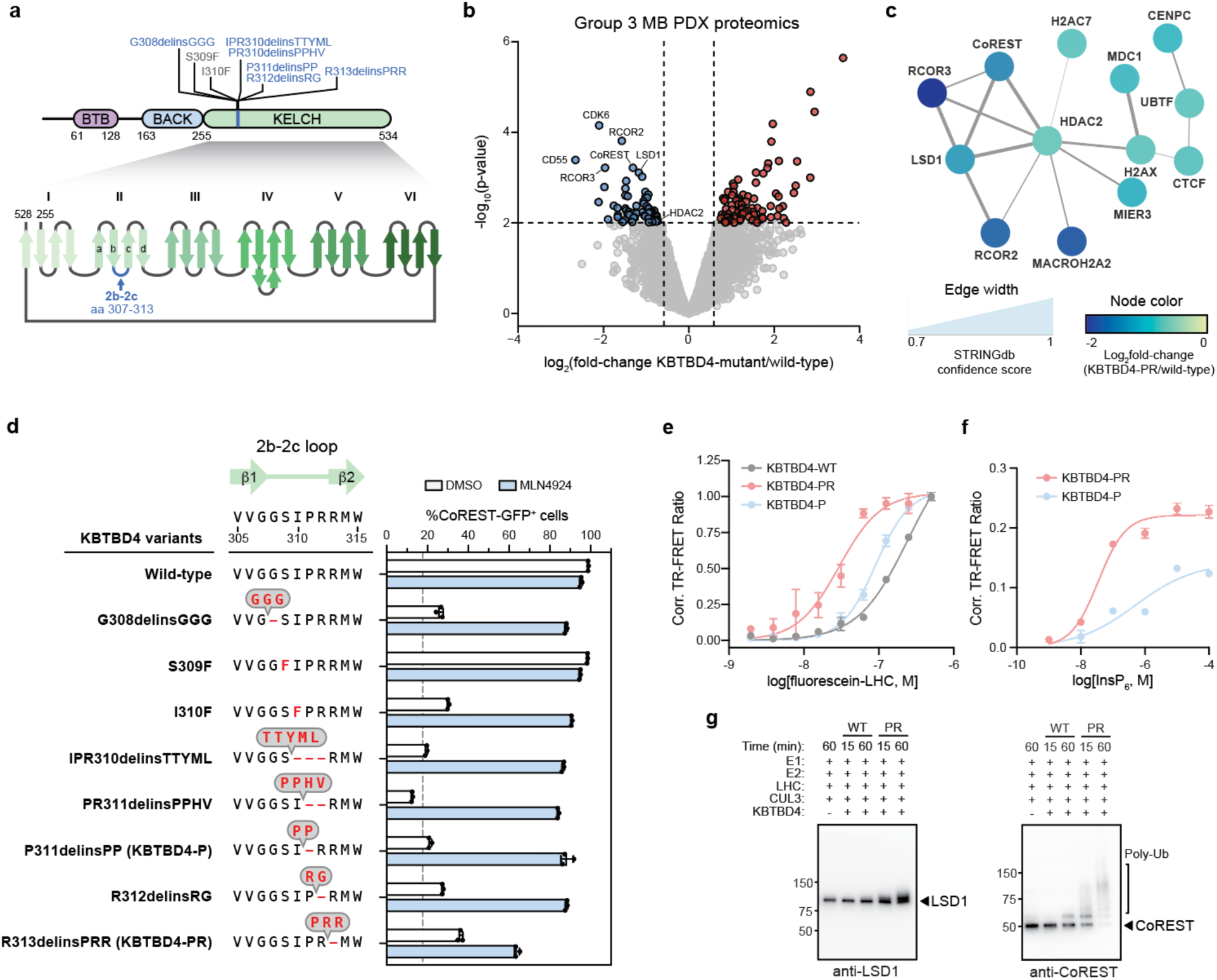
KBTBD4 MB mutants potentiate E3 activity. a) Schematic of the protein domains of KBTBD4 (top) and recurrent mutations in MB at a hotspot in the KELCH domain blade II (bottom). b) Volcano plot showing proteins up- and down-regulated in *KBTBD4^MUT^* (*n =* 2) versus WT (*n =* 5) PDX models. Colored dots (blue, red) show proteins with | log_2_(fold-change) | > 0.7 in *KBTBD4^MUT^* versus WT and p-value < 0.01. Red and blue dots depict proteins that are up- and down-regulated, respectively. c) Protein STRING network showing proteins significantly depleted in *KBTBD4^MUT^* (*n =* 2) versus WT (*n =* 5) PDX models. Node color scale depicts log_2_(fold-change) protein abundance in mutant versus WT models. Edges indicate PPIs, and the width scale depicts confidence of the PPI for the nodes. d) Flow cytometry quantification of GFP^+^ cells (%, *x*-axis) for KBTBD4-null CoREST-GFP cells overexpressing indicated KBTBD4 variant and treated with either DMSO or MLN4924 (1 µM) for 24 h. Bars represent the mean ± s.d. of *n =* 3 replicates, dots show the individual replicate values. e) Dose-response curve showing TR-FRET signal (*y*-axis) between fluorescein-LHC (*x-*axis) and anti-His CoraFluor-1-labelled antibody with indicated His-KBTBD4 variant in the presence of InsP_6_ (50 µM). Data represent the mean ± s.d. of *n =* 2 replicates. f) Dose-response curve showing TR-FRET signal (*y*-axis) between fluorescein-LHC and anti-His CoraFluor-1-labelled antibody with indicated His-KBTBD4 mutant in the presence of varying concentrations of InsP_6_ (*x*-axis). Data represent the mean ± s.d. of *n =* 2 replicates. g) Immunoblots of in vitro ubiquitination assays of CRL3^KBTBD4-WT^ and CRL3^KBTBD4-PR^ with LHC. Results in **d-g** are representative of two independent experiments.

### KBTBD4 MB mutants potentiate E3 activity and CoREST degradation

To determine how *KBTBD4* mutations might remodel the MB proteome, we conducted total proteomic profiling in patient-derived xenograft (PDX) models derived from Group 3 MB tumor specimens, including two harboring the *KBTBD4*-PR mutation and five *KBTBD4-*wild-type (WT). Comparison of the WT and PR-mutant models identified 64 and 82 proteins that were significantly up- and down-regulated, respectively, in the mutant samples (**Fig. 1b**). Functional network analysis of the differentially expressed proteins revealed that a network of HDAC2- associated proteins was highly depleted (**Fig. 1c; Extended Data Fig. 1a, b**). Notably, this included members of the CoREST corepressor complex (i.e., RCOR1 (CoREST), RCOR2, RCOR3, LSD1) as well as MIER3, another HDAC1/2-associated corepressor. These results are consistent with our prior data showing that overexpression of KBTBD4-P, KBTBD4-PR, and R312delinsRG, in MB cell lines induced LSD1-CoREST degradation^4^. Altogether, these results demonstrate that several HDAC1/2-associated corepressors, most notably LHC, are selectively depleted in clinically relevant KBTBD4-mutant PDX models of Group 3 MB.

The absolute and relative activities of other KBTBD4 MB mutations remain unknown. Consequently, we measured the ability of all known MB mutations to affect CoREST stability. Using a K562 cell line containing GFP knocked in-frame at the CoREST C-terminus as well as KBTBD4 knockout (CoREST-GFP/KBTBD4-null cells), we confirmed that overexpression of KBTBD4-P and KBTBD4-PR, but not WT ligase, were sufficient to induce CoREST-GFP depletion (**Fig. 1d; Extended Data Fig. 2a-c**). Except for S309F, all other known MB mutations could significantly enhance CoREST-GFP depletion. Enhanced CoREST-GFP degradation could be largely blocked by addition of MLN4924, a neddylation inhibitor that disrupts the activity of CULLIN-RING E3 ligase systems. Collectively, these results show that diverse KBTBD4 MB mutations can promote CoREST degradation.

We next sought to establish that the KBTBD4-mutant E3 ligase directly engages and ubiquitinates the CoREST corepressor complex using a reconstituted system. We started with the biochemically purified LHC complex containing HDAC1 — an interchangeable paralog of HDAC2. Within this CoREST core complex, CoREST (aa 84-482) was labeled with fluorescein (see Methods)^22,23^. We also purified His-KBTBD4, His-KBTBD4-P, and His-KBTBD4-PR. With reconstituted complexes in hand, we measured their association in vitro using time-resolved Förster resonance energy transfer (TR-FRET)^24,25^ (**Fig. 1e**). Remarkably, His-KBTBD4-P and His- KBTBD4-PR both demonstrated greater affinity with LHC than WT, showing that the MB mutants are sufficient to drive E3 engagement with CoREST in the context of the LHC core. KBTBD4-PR exhibited significantly stronger binding to LHC than KBTBD4-P, which was recapitulated by co-immunoprecipitation (co-IP) experiments (**Extended Data Fig. 1c**). Critically, mutant KBTBD4- LHC binding required the addition of inositol hexakisphosphate (InsP_6_) (**Fig. 1f**), a cofactor that stabilizes the protein-protein interaction between CoREST and HDAC1^26^. This dependency on InsP_6_ further supports the necessity of HDAC1/2 in the mechanism of mutant KBTBD4. Lastly, reconstituted CRL3^KBTBD4-PR^ exhibited increased ubiquitination of LHC in vitro in comparison to CRL3^KBTBD4-WT^ (**Fig. 1g; Extended Data Fig. 1d, e**). Together, these data demonstrate that MB mutations in the 2b-2c loop of KBTBD4 are sufficient to increase ubiquitination and degradation of CoREST complexes in an HDAC1/2-dependent fashion.

### Deep mutational scanning of the 2b-2c loop

Point substitutions and indels can incur fundamentally different impacts on protein structure and function, with the latter typically having more severe effects^27–29^. In the case of KBTBD4, we showed that a point substitution and diverse indels can both enhance CoREST degradation, yet indels in KBTBD4 occur most frequently in patients. The bias of KBTBD4 MB mutations toward indels is not understood, possibly reflecting their increased likelihood in promoting gain-of-function interactions or the error-prone processes (i.e., copy count variants from faulty DNA replication) that favor their formation^30,31^.

To address these possibilities, we used deep mutational scanning to profile the mutational landscape of the 2b-2c loop and systematically compare the effects of point substitutions and indels on CoREST degradation. Specifically, we constructed a library of KBTBD4 mutants (5,220 total) comprising all possible 1 amino acid (aa) deletions (5), 1 aa substitutions (133), 2 aa substitutions involving pairs of adjacent residues (2,166), 1 aa insertions (134), and 2 aa insertions (2,680) across the 7 aa sequence spanning Gly307 and Arg313; as well as 100 randomly scrambled WT sequences and the 2 remaining MB indels (PR311delinsPPHV, IPR310delinsTTYML) not encompassed in the aforementioned categories (**Fig. 2a**). This mutant pool was transduced into CoREST-GFP/KBTBD4-null cells and after 3 days, GFP^−^ and GFP^+^ cells were sorted by FACS. The identities of enriched and depleted KBTBD4 variants, respectively, were then determined using next-generation sequencing and compared to variant frequencies from transduced, unsorted cells (**Fig. 2b; Extended Data Fig. 2c**).

**Figure 2.**
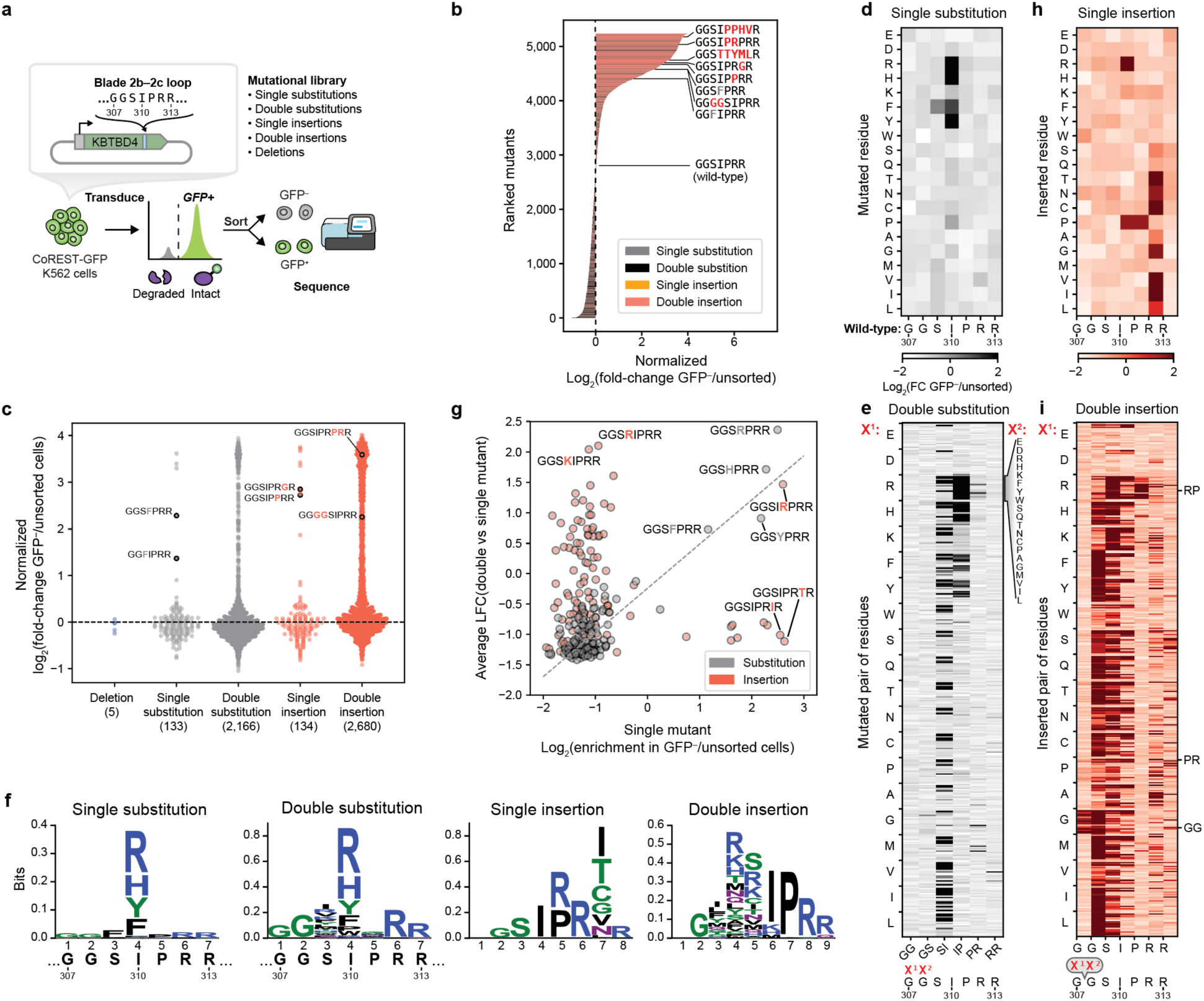
Deep mutational scanning of the 2b-2c loop. a) Schematic of the KBTBD4 2b-2c loop deep mutational scanning. b) Waterfall plot displaying KBTBD4 variants ranked (*y*-axis) by their log_2_ fold-change enrichment in GFP^−^ over unsorted population normalized to WT (*x*-axis). Bars represent mean of *n* = 3 replicates. The color scheme represents single substitution (dark gray), double substitution (black), single insertion (orange), double insertion (red) and miscellaneous (light gray). WT and MB mutant sequences are labeled on the plot. c) Swarm plot showing fold-change enrichment of KBTBD4 variants in GFP^−^ cells normalized to WT (*y-*axis) classified by mutation type. Dotted line indicates fold-change enrichment of WT KBTBD4, and MB mutant sequences are labeled. d) Single-substitution deep mutational scanning displayed as a heatmap of fold-change enrichment in GFP^−^ cells for each mutated amino acid. e) Double-substitution deep mutational scanning displayed as a heatmap of fold-change enrichment in GFP^−^ cells for each mutated amino acid pair. The *x*-axis indicates the positions of the mutated amino acid pairs (X^1^X^2^), with the identities of the substituted residues shown on the *y*-axis. Specifically, the first substituted residue (X^1^) is indicated by the lefthand labels, while the second residue (X^2^) is indicated by each row. f) Sequence logo depicting relative entropy of amino acids at each position for single substitution, double substitution, single insertion, and double insertion mutant sequences. One letter codes of amino acids are colored by their chemical characteristics: hydrophobic (black), polar (green), basic (blue), acidic (red), and neutral (purple). g) Scatterplot showing fold-change enrichment of single mutant KBTBD4 variants at the Nth position (either substitution or insertion) in GFP^−^ cells (*x* axis) and average fold-change of the corresponding double mutants created by mutation of the adjacent N-1 or N+1 position. Substitutions and insertion mutants are colored in gray and red, respectively, and selected mutant sequences are labeled. Linear correlation (dotted line) based on linear least-squares regression for the substitutions are displayed on the plot (Pearson correlation coefficient *r* = 0.856, p-value = 1.74 × 10^-39^). h) Single-insertion deep mutational scanning displayed as a heatmap of fold-change enrichment in GFP^−^ cells for each mutated amino acid. i) Double-insertion deep mutational scanning displayed as a heatmap of fold-change enrichment in GFP^−^ cells for each inserted amino acid pair. The *x*-axis indicates the positions of the inserted amino acid pairs (X^1^X^2^), with the identities of the inserted residues shown on the *y*-axis. Specifically, the first inserted residue (X^1^) is indicated by the lefthand labels, while the second residue (X^2^) is indicated by each row. Data in **b-i** are mean of *n =* 3 replicates and the overall deep mutational scanning experiment was performed once.

As anticipated, KBTBD4 variants enriched in GFP^−^ cells (i.e., CoREST-GFP degraded) were strongly depleted in GFP^+^ cells (i.e., CoREST-GFP intact) (**Extended Data Fig. 2d**). Reassuringly, the WT KBTBD4 sequence was depleted in GFP^−^ cells, while the MB mutations were enriched to varying degrees, with the insertion MB mutants generally outperforming the MB point substitutions. While deletions had no impact on CoREST-GFP degradation, both substitutions and insertions could significantly promote degradation — with the double amino acid perturbations generally outperforming the single amino acid perturbations (**Fig. 2c**). In particular, the double insertions as a class were the most effective at promoting CoREST-GFP degradation overall (**Fig. 2b, c**), in full agreement with their biased enrichment as MB mutations.

We next scrutinized the amino acid backbone positions and side chain alterations within each mutant category that most effectively enhanced CoREST-GFP degradation. Efficacious single substitutions highly favored mutations at position 4 (i.e., Ile310) to bulkier positively charged (Arg, His) or aromatic amino acids (Phe, Tyr), including the I310F MB mutant (**Fig. 2c, d**). This positional and amino acid preference was maintained for the double substitutions, where mutation of Ile310 (i.e., second position of SI or first position of IP) was highly favored (**Fig. 2e, f; Extended Data Fig. 3a**). In fact, the most effective double substitutions were derived from the most effective single substitutions by further mutation of an adjacent neighboring position (i.e., Ser309 or Pro311) (**Fig. 2g**, Pearson’s *r* = 0.856, p-value = 1.74 × 10^-39^). By contrast, efficacious single insertions showed less positional and amino acid bias (**Fig. 2h, f**), albeit insertions after position 6 (i.e, Arg312) were favored. Amino acids inserted after position 6 could be diverse, suggesting that the primary function of the insertion may be to shift the 2b-2c loop so that Arg312 is moved to position 5. Supporting this notion, insertion of an Arg residue after position 4 (i.e., Ile310) was uniquely potent for CoREST-GFP degradation, which, together with our single substitution data, suggests that introduction of an Arg residue in the middle of the loop is heavily favored.

Strikingly, effective double insertions significantly diverged from the single insertions, showing stronger preference for insertion earlier in the 2b-2c loop sequence (**Fig. 2f; Extended Data Fig. 3b**). This positional preference was less pronounced when Arg, and to a lesser extent Lys, Met, and Pro, constituted one of the inserted amino acids. Notably, effective double insertions could tolerate many more types of amino acid side chains, except acidic residues (Asp, Glu) or Trp. These relaxed preferences likely explain the enhanced performance of double insertions in the deep mutational scan (**Fig. 2c**), suggesting they may operate to introduce a bulkier or basic amino acid into the loop by either shifting the position of Ile310 or through direct insertion of such a residue. Notably, in contrast to the relationship between single and double substitutions, the best double insertions were generally not derived from the most effective single insertions (i.e., by insertion of an adjacent residue) (**Fig. 2g**, Pearson’s *r* = –0.04, n.s.). These data suggest that (1) single and double insertions remodel the 2b-2c loop by distinct mechanisms, in contrast to single and double substitutions, and that (2) insertions are particularly effective in promoting gain-of-function protein-protein interactions^28^. Altogether, our deep mutational scanning supports the notion that double insertions are enriched in KBTDB4-mutant MBs due to their propensity to promote neomorphic activity.

### Overall structures of LHC-bound KBTBD4 mutants

Our deep mutational scanning highlighted the potency of double insertions in promoting KBTBD4 gain-of-function. To elucidate the mechanism by which double insertions enable high-affinity engagement of the LHC complex, we determined the cryo-EM structures of two LHC- bound KBTBD4 MB mutants, KBTBD4-PR and KBTBD4-TTYML. We chose the KBTBD4-TTYML mutant in addition to the most common KBTBD4-PR mutant because it lacks a basic residue in the 2b-2c loop. The KBTBD4 dimer is well resolved in both structures, which were determined at 3.84 and 3.30 Å resolution for the PR and TTYML mutants, respectively (**Fig. 3a-c**; **Extended Data Fig. 4, 5, Extended Data Table 1**). The 3D reconstruction maps enabled us to trace the entire E3 polypeptide in both structures with high confidence.

**Figure 3.**
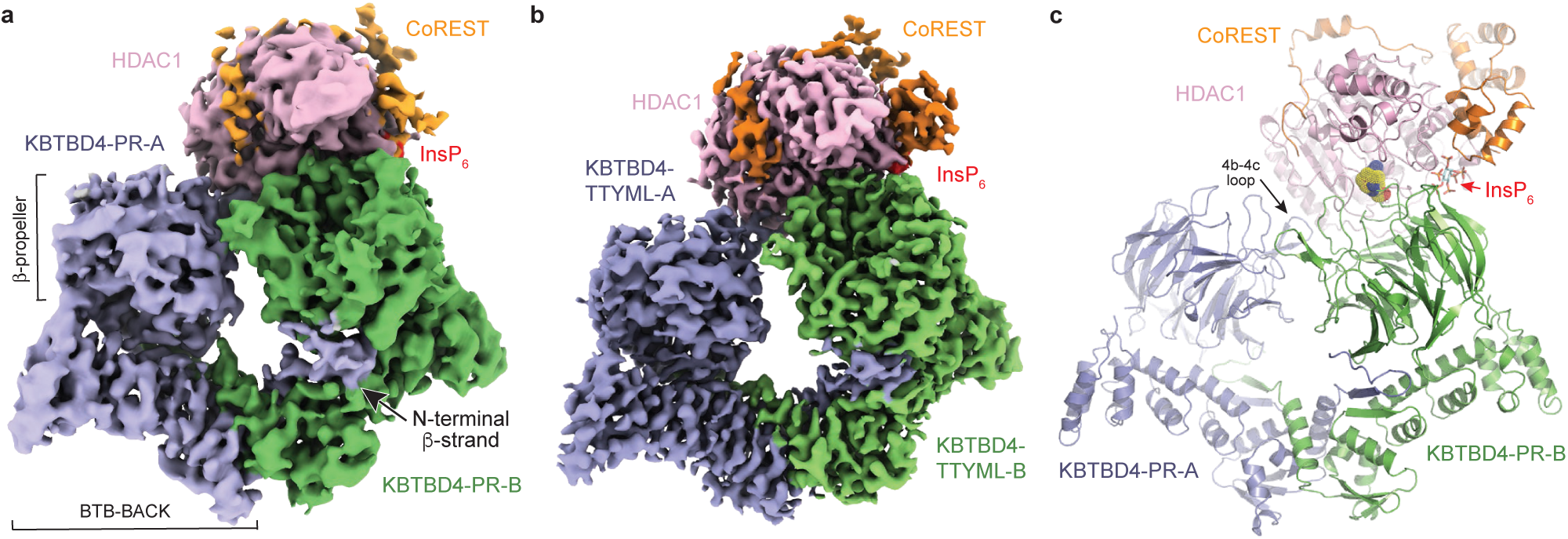
Overall structures of LHC-bound KBTBD4 mutants. a) Cryo-EM map of LHC-bound KBTBD4-PR mutant with the two KBTBD4-PR protomers colored in slate and green, HDAC1 colored in pink, CoREST colored in orange, and InsP_6_ colored in red. b) Cryo-EM map of LHC-bound KBTBD4-TTYML mutant. The color scheme is the same as in **a**. c) Ribbon diagram of the KBTBD4-PR-HDAC1-CoREST-InsP_6_ complex. The subunits of the complex are colored the same way as in **a.** The hot spot arginine residue is shown in space filling model mode. InsP_6_ is shown in cyan and red sticks.

As expected, the KBTBD4 protein adopts a well-defined dimeric architecture with a pseudo-two-fold symmetry (**Fig. 3c**). In a head-to-head fashion, both KBTBD4 mutants homodimerize via its N-terminal BTB domain with its C-terminal KELCH-repeat domain positioned atop the elongated BTB-BACK structure module. Despite their apparent two-fold symmetry, the two mutant KBTBD4 dimers bind LHC in an asymmetric, concerted assembly (**Fig. 3a-c**). In both cryo-EM maps, only HDAC1 and part of the ELM2 and SANT1 domains of CoREST are visible, although the two CoREST domains are better resolved in the KBTBD4-TTYML-LHC complex. The rest of the LHC complex is presumably too flexible to be visualized by 3D reconstruction.

Astonishingly, HDAC1 is the only subunit of the LHC complex that makes direct contact with the E3 mutants. In both E3-substrate assemblies, the KBTBD4 dimer engages LHC by asymmetrically cupping the catalytic domain of HDAC1 through its two β-propellers. Unexpectedly, in contrast to other KELCH CRL3s, the mutant E3 dimers do not use the top surface of their β-propellers to recognize HDAC1. In the KBTBD4-PR mutant dimer, one E3 protomer, designated as KBTBD4-PR-A, uses the 4b-4c loop in its β-propeller to latch onto the edge of the HDAC1 catalytic domain, whereas the other protomer, KBTBD4-PR-B, makes more extensive contact with the enzyme by wrapping around its active site via a lateral surface region of its KELCH-repeat domain (**Fig. 3c**). These HDAC1-binding regions of the two E3 protomers are distinct and positioned away from the KBTBD4-HDAC1 interface in the opposite protomer. Notably, next to the HDAC1 active site, the interface between the deacetylase and KBTBD4-PR- B is stabilized by InsP_6_ (**Fig. 3c**). By making direct contacts with HDAC1, CoREST, and KBTBD4, InsP_6_ acts as an essential molecular glue at the protein interface (**Fig. 1d**). The two β-propellers of KBTBD4, therefore, use distinct structural elements to engage the deacetylase.

### Structural mechanisms of KBTBD4 MB gain-of-function mutations

Superposition analysis of the two KBTBD4-PR protomers indicates that the relative position of the β-propeller domain and the BTB-BACK domain is not identical in the two E3 chains (**Fig. 4a**). The position of each of the two β-propeller domains relative to the dimeric BTB-BACK platform is most likely optimized for binding HDAC1, consistent with a global induced-fit mechanism for assembling the E3-substrate complex. Similar to the BTB-BACK module, the KELCH-repeat domain of each KBTBD4 mutant maintains the same structure throughout the entire β-propeller fold except at a few surface loop regions (**Extended Data Fig. 6a**). In particular, the 4b-4c loop and the 2b-2c loop, which are uniquely employed by propeller-A and -B to interact with HDAC1, respectively, show clear structural deviations between the two protomers (**Fig. 4b- d**). In KBTBD4-TTYML, the mutant 2b-2c loop in propeller-A is solvent-exposed and structurally disordered. By contrast, the same loop in propeller-B adopts an ordered conformation, making close interactions with HDAC1 (**Fig. 4d**). Overall, the E3 dimer demonstrates both global and local structural plasticity to engage the neo-substrate.

**Figure 4.**
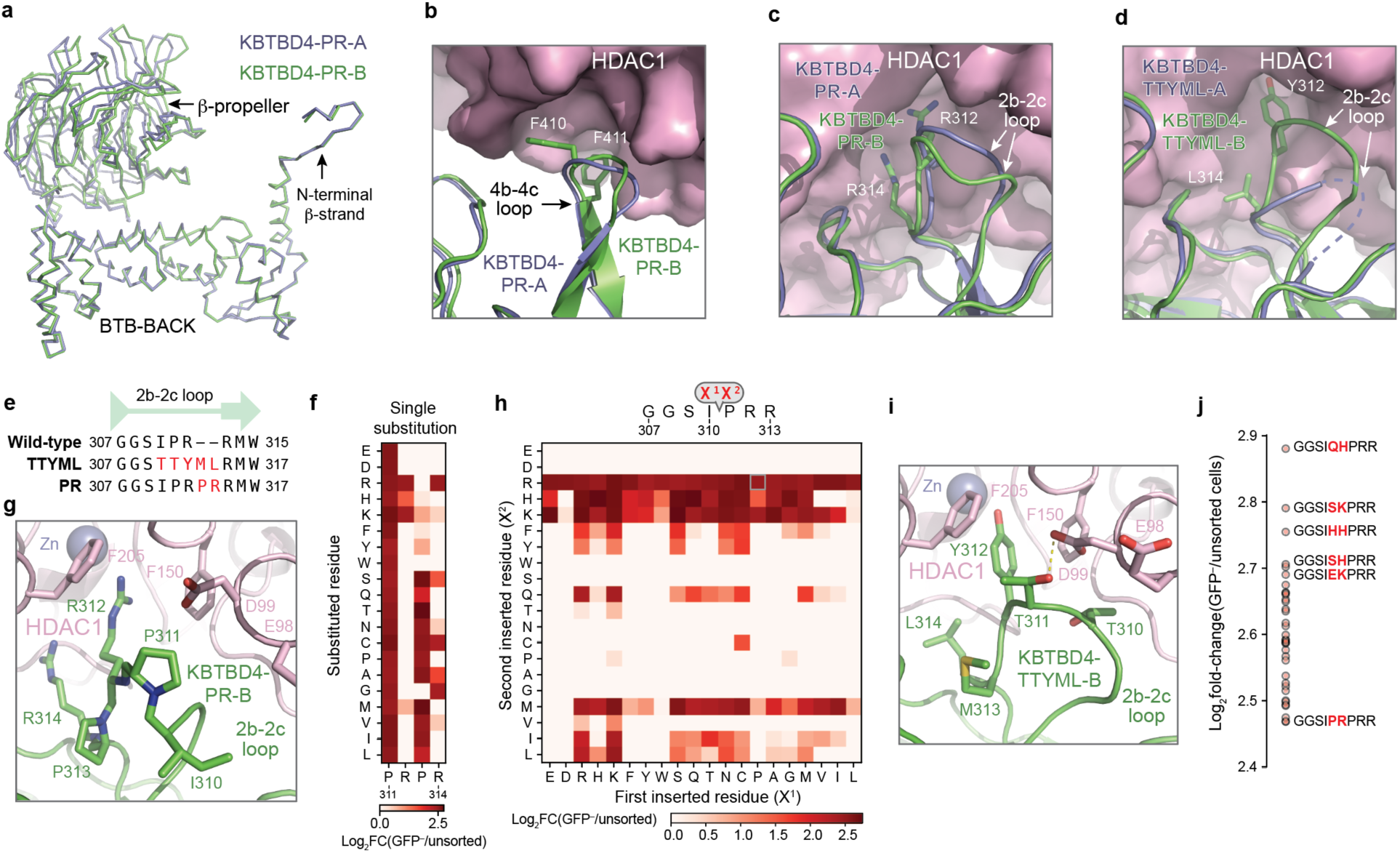
Structural mechanisms and amino acid preferences of functional KBTBD4 mutations. a) Superposition of the two KBTBD4-PR protomers in complex with HDAC1. The two protomers, KBTBD4-PR-A (slate) and KBTBD4-PR-B (green) are superimposed via their BTB-BACK domain. b) A close-up view of the 4b-4c loops of KBTBD4-PR-A (slate) and KBTBD-PR-B (green) after the β-propeller domain of the former is superimposed onto that of the latter. The side chains of two phenylalanine residues in the 4b-4c loop of KBTBD4-PR-B are shown in sticks. c) A close-up view of the 2b-2c loops of KBTBD4-PR-A (slate) and KBTBD-PR-B (green) after the β-propeller domain of the former is superimposed onto that of the latter. The side chains of two arginine residues in the 2b-2c loop of KBTBD4-PR-B are shown in sticks. d) A close-up view of the 2b-2c loops of KBTBD4-TTYML-A (slate) and KBTBD-TTYML-B (green) after the β-propeller domain of the former is superimposed onto that of the latter. The side chains of two interface residues in the 2b-2c loop of KBTBD4-PR-B are shown in sticks. e) Alignment of KBTBD4 WT, PR, and TTYML 2b-2c loop sequences. f) Single-substitution deep mutational scanning (i.e., saturation mutagenesis) displayed as a heatmap of fold-change enrichment in GFP^−^ cells for each mutated amino acid in the KBTBD4- PR PRPR sequence. Color intensity represents mean of *n* = 3 replicates. g) A close-up view of the interface between HDAC1 (pink) and the 2b-2c loop of KBTBD4-PR-B (green) with the side chains of key residues shown in stick. h) Double-insertion deep mutational scanning displayed as a heatmap of fold-change enrichment in GFP^−^ cells for each pair of mutated amino acids (X^1^, X^2^) inserted after Ile310. Color intensity represents mean of *n* = 3 replicates. i) A close-up view of the interface between HDAC1 (pink) and the 2b-2c loop of KBTBD4-PR-B (green) with the side chains of key residues shown in stick. j) Top-enriched double-insertion mutants inserted after Ile310, more effective than KBTBD4- PR, ranked by their log_2_ fold-change enrichment in GFP^-^ over unsorted population (*y*-axis). Dots represent mean of *n* = 3 replicates. Data in **f**, **h**, and **j** are mean of *n =* 3 replicates and the overall deep mutational scanning experiment was performed once.

The indel mutations in the PR and TTYML mutants elicit their gain-of-function effects at the center of the HDAC1 propeller-B interface, which represents the hotspot protein-protein interaction site. The net effects of these two mutations are the expansion of the 2b-2c loop by two amino acids and, in the case of TTYML, alteration of the amino acid composition (**Fig. 4e**). Surrounded by nearby structural elements that also make direct contacts with HDAC1, the 2b-2c loops of the two KBTBD4 mutants are positioned right at the rim of the active site pocket of the deacetylase. Despite their sequence diversity, both mutants insert the bulky side chain of a central residue into the narrow tunnel leading to the catalytic site of the enzyme. For KBTBD4-PR, the positively charged side chain of Arg312 reaches halfway into the tunnel and is accompanied by Arg314, which occupies a nearby outer vestibule at the tunnel’s entrance (**Fig. 4c**). In an analogous manner, the aromatic side chain of Tyr312 in KBTBD4-TTYML also protrudes into the HDAC1 tunnel and is supported by Leu314, which partially fulfills the role of Arg314 in the KBTBD4-PR mutant (**Fig. 4d**).

Taking advantage of our deep mutational scanning data, we next investigated the requirement of each residue in the central PRPR sequence of KBTBD4-PR for CoREST degradation (**Fig. 4f; Extended Data Fig. 7**). A basic residue is strongly preferred at the second position in this motif, corroborating the critical role played by Arg312 at the interface (**Fig. 4g**). Arg314, however, can be replaced by amino acids with a smaller side chain, suggesting an auxiliary function of the residue. By contrast, the amino acid preferences at the two proline positions are more relaxed, which is consistent with their minor involvement in contacting HDAC1. However, if the first proline of the PRPR motif is replaced by a polar residue (X^1^ position), such as serine or lysine, the requirement for a basic amino acid at the X^2^ position becomes less stringent (**Fig. 4h**). In this context, a hydrophobic or aromatic amino acid can now substitute the basic residue to effectively secure HDAC1, presumably by inserting its bulky side chain into the active site tunnel of the enzyme. In fact, this notion is validated by the TTYML mutant, in which Tyr312 of the central TTYML motif plays a role equivalent to Arg312 in the PRPR motif (**Fig. 4i**). Although tyrosine is not as strongly favored as arginine (**Fig. 4h**), the two leading threonine residues, Thr310 and Thr311, of the TTYML mutant most likely compensate by making closer contacts with an HDAC1 surface loop — donating a hydrogen bond from Thr310 to Asp99 of HDAC1 (**Fig. 4i**). These additional interactions outside the HDAC1 active site tunnel provides a plausible explanation for the relaxed requirement at the second position of the PRPR motif when the first proline is replaced by a polar residue (**Fig. 4h**) and the superior activity of QHPR, SKPR, and HHPR over PRPR (**Fig. 4j**). These synthetic mutants and the cancer mutants, therefore, exploit not only the active site tunnel of HDAC1, but also its peripheral regions. Taken together, we conclude that the KBTBD4 gain-of-function mutants enable the productive interaction between the E3 and the neo-substrate by augmenting their shape complementarity and polar interactions.

### Converging mechanism between KBTBD4 MB mutations and UM171

Recent studies have shown that KBTBD4 is involved in the mechanism of action of UM171, a potent agonist of *ex vivo* hematopoietic stem cell expansion (**Fig. 5a**)^21^. Remarkably, UM171 phenocopies the KBTBD4 MB mutations by promoting the ubiquitination and degradation of CoREST. In a companion study, we determined the structure of the KBTBD4-LHC complex stabilized by UM171 and demonstrated that the small molecule acts as a molecular glue to induce complex formation between the E3 and HDAC1. A structural comparison between the KBTBD4- UM171-LHC complex and the two LHC-bound KBTBD4 MB mutants unveils a striking converging mechanism by which the molecular glue degrader and the cancer mutations complement and optimize the suboptimal protein-protein interface between the E3 and HDAC1 to drive their association.

**Figure 5.**
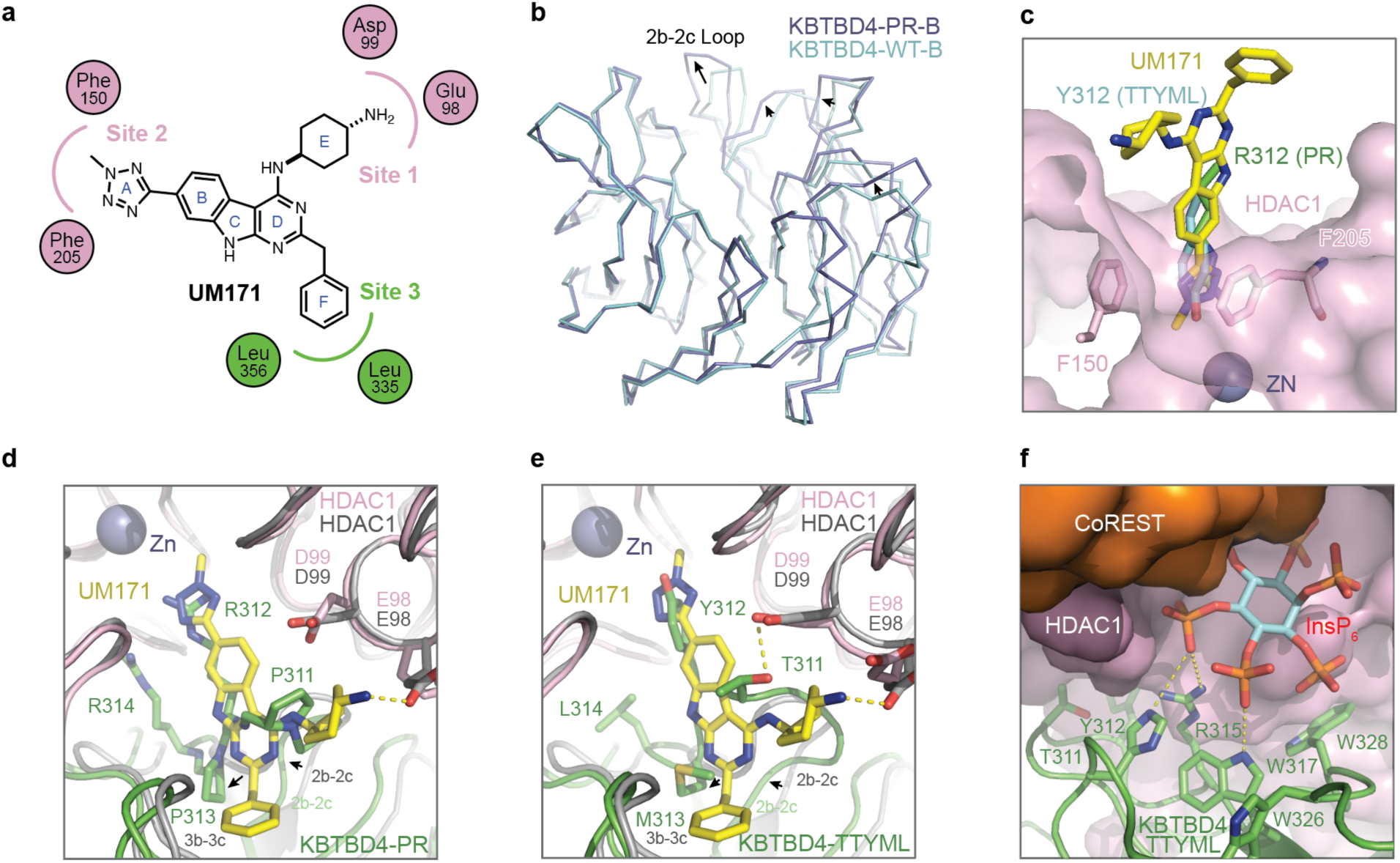
Converging mechanism between KBTBD4 cancer mutations and UM171. a) Simplified ligand plot of UM171-KBTBD4-HDAC1 interactions. HDAC1 and KBTBD4 residues are denoted by pink and green circles, respectively. b) Superposition analysis of the β-propellers in protomer-B of the KBTBD4-WT and KBTBD4-PR dimers. The structural differences at several top surface loops are indicated by arrows. 2b-2c loop is labeled. c) A comparison of UM171 (yellow and blue sticks), the side chain of Tyr312 of KBTBD4-TTYML- B (cyan and red sticks), and the side chain of Arg312 of KBTBD4-PR-B (green and blue sticks) at the active site pocket of HDAC1 (pink). The three complex structures are superimposed via HDAC1. ZN: zinc. Two phenylalanine residues outlining the entrance of the HDAC1 active site tunnel are shown in sticks. d) A comparison between UM171 (yellow and blue sticks) and the 2b-2c loop of KBTBD4-PR-B with the KBTBD4-UM171-HDAC1 structure superimposed with the KBTBD4-PR-HDAC1 structure via HDAC1. The side chains of key residues at the interface are shown in sticks. e) A comparison between UM171 (yellow and blue sticks) and the 2b-2c loop of KBTBD4- TTYML-B with the KBTBD4-UM171-HDAC1 structure superimposed with the KBTBD4- TTYML-HDAC1 structure via HDAC1. The side chains of key residues at the interface are shown in sticks. f) A close-up view of the inter-molecular interfaces among KBTBD4-TTYML (green), HDAC1 (pink, surface representation), CoREST (orange, surface representation), and InsP_6_ (cyan, orange, and red sticks). The side chains of key KBTBD4-TTYML residues involved in InsP_6_ interaction and at the nearby 2b-2c loop are shown in sticks.

As expected, the double insertion of the KBTBD4-PR mutant expands the 2b-2c loop and triggers subsequent conformational changes in a series of additional top surface loops across half of the β-propeller (**Fig. 5b**). Due to these changes, the relative position of HDAC1-CoREST is slightly shifted when the KBTBD4-UM171-LHC complex is superimposed with the KBTBD4-PR- LHC complex via propeller-B (**Extended Data Fig. 6b**). Nonetheless, global superposition of the two complex structures can be made with an R.M.S.D. of 1.0 Å over ∼1,500 Cα atoms, indicative of a highly similar overall architecture (**Extended Data Fig. 6c**). Indeed, the two E3-substrate assemblies are stabilized by the same protein-protein interfaces overall and share the common hotspot interactions centered around the active site pocket of HDAC1.

Closer inspection of the active site hotspot reveals that UM171 acts as a chemical facsimile of the KBTBD4 MB mutations. Simplistically, UM171 possesses three ‘arms’, a cyclohexylamine (E ring), an *N-*methyltetrazole (A ring), and a benzyl group (F ring), which roughly occupy three sites that are also contacted by three distinct amino acids of mutant KBTBD4 (**Fig. 5a**). First, near the entrance of the pocket, the cyclohexylamine of UM171 and the X^1^ position of the two cancer mutants (i.e., Pro311 of KBTBD4-PR and Thr311 of KBTBD4-TTYML) both contact the same HDAC1 loop comprising Asp99 and Glu98 (**Fig. 5d, e**, site 1). Second, the tetrazole of UM171 mimics the side chain of the central amino acid at the mutants’ X^2^ position (e.g., Arg312 of KBTBD4-PR and Tyr312 of KBTBD4-TTYML), where both protrude into the HDAC1 active site tunnel, reaching the same depth (**Fig. 5c**, site 2). Lastly, while the benzyl group of UM171 induces and binds a surface groove between the 2b-2c and 3b-3c loops of KBTBD4 (site 3), the expanded 2b-2c loops of the two cancer mutants (i.e., Pro313 of KBTBD4-PR and Met313 of KBTBD4- TTYML) occupy the same space — enabling them to project their central bulky amino acid into the HDAC1 active site (**Fig. 5d, e**). Importantly, the structural roles of UM171 and the double- insertions in the two KBTBD4 mutants are mostly localized at the active site of HDAC1, allowing InsP_6_ to further strengthen the E3-substrate interaction through a separate contact interface (**Fig. 5f**). Therefore, the molecular glue compound and the gain-of-function cancer mutations physically and functionally mimic each other, complementing the suboptimal protein-protein interface to promote the neomorphic interaction.

### HDAC1/2 inhibitors block mutant KBTBD4 neomorphic activity

Our cryo-EM structure reveals that MB mutations in KBTBD4 promote HDAC1/2 engagement by inserting a residue into the deacetylase active site, further extending the protein-protein interaction interface between the E3 and neo-substrate. Our deep mutational scanning shows the requirement of this hotspot interaction within the HDAC1/2 active site for CoREST degradation. Together, these observations suggest that active site HDAC1/2 inhibitors may disengage KBTBD4 mutants by sterically occluding the mutant ligase, thereby blocking their oncogenic function. Superposition analysis of KBTBD4-PR-HDAC1 with HDAC2 bound to suberoylanilide hydroxamic acid (SAHA, also known as vorinostat) supports the notion that the small molecule and inserted arginine residue might physically clash (**Fig. 6a**)^32^.

**Figure 6.**
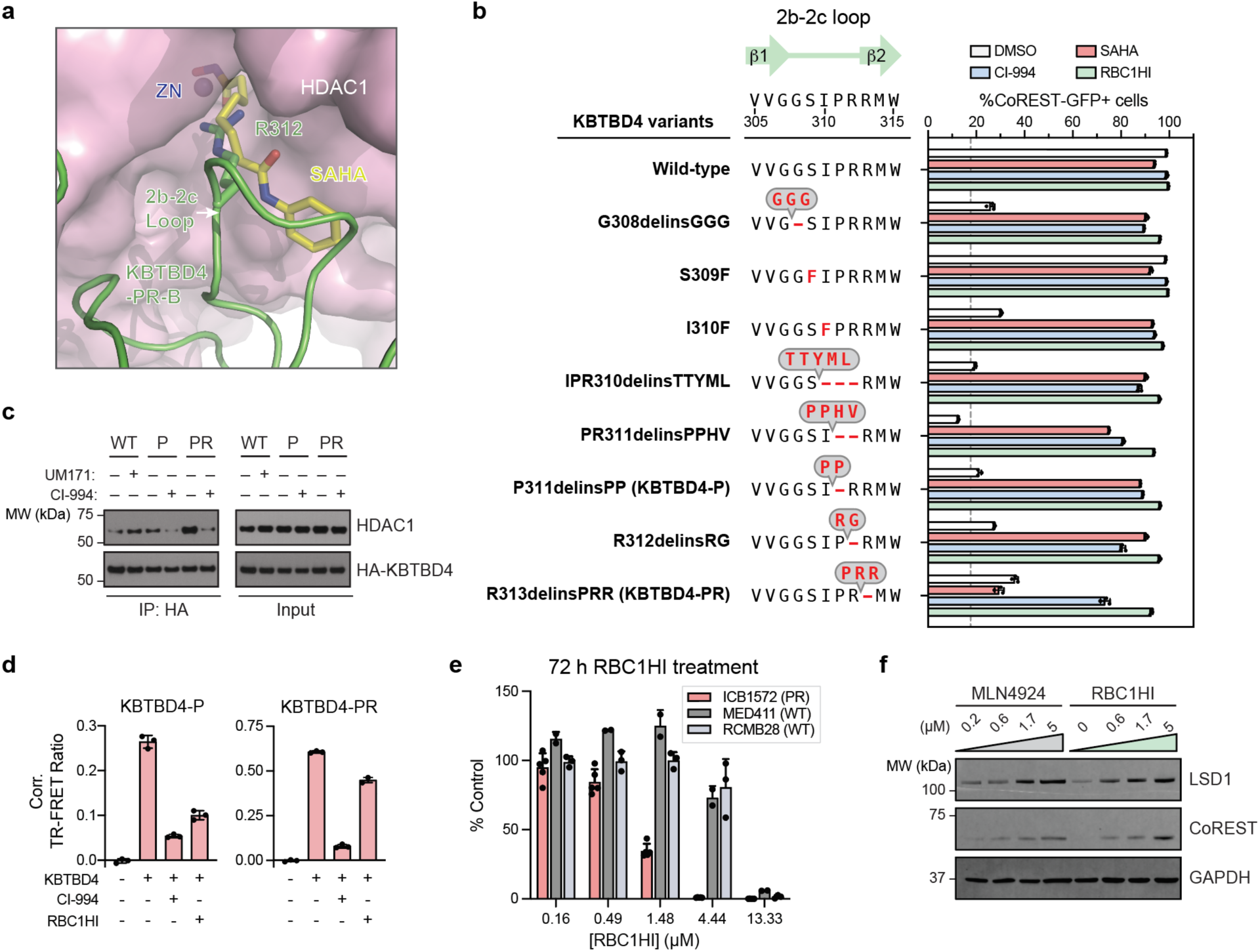
HDAC1/2 inhibitors block the neomorphic activity of KBTBD4 mutants. a) Steric clash between SAHA and the central arginine residue at the 2b-2c loop of KBTBD4- PR-B. The KBTBD4-PR-HDAC1 complex structure is superimposed with the HDAC2-SAHA complex structure (PDB:4LXZ) via the HDAC subunits. b) Flow cytometry quantification of GFP^+^ cells (%, *x*-axis) for KBTBD4-null CoREST-GFP cells overexpressing indicated KBTBD4 variant and treated with either DMSO, CI-994 (10 µM), SAHA (10 µM), or RBC1HI (10 µM) for 24 h. Bars represent the mean ± s.d. of *n* = *3* replicates, dots show the individual replicate values. c) Immunoblots of HA IP in the presence of DMSO, UM171 (1 µM, 1 h), or CI-994 (10 µM, 3 h), and MLN4924 (1 µM, 3 h) from 293T cells transfected with indicated HA-KBTBD4 variant. d) TR-FRET signal (*y*-axis) between fluorescein-LHC and anti-His CoraFluor-1-labeled antibody with indicated His-KBTBD4 mutant in the presence of DMSO, CI-994 (10 µM), or RBC1HI (10 µM). Bars represent the mean ± s.d. of *n* = 3 replicates, dots show the individual replicate values. e) Proliferation assay for ICB1572 (*KBTBD4-PR*) and MED411 (WT *KBTBD4*) ex vivo cell proliferation (*y*-axis, % control) after RBC1HI treatment for 72 h at indicated doses (*x*-axis). Data represent mean ± s.d. across cells from *n* = 5 (ICB1572), *n* = 3 (RCMB28), and *n* = 2 (MED411) mice. f) Immunoblot showing LSD1, CoREST, and GAPDH in ICB1572 (*KBTBD4-PR*) after 24 h treatment with MLN4924 or RBC1HI at the indicated doses. Results in **b-d, e** are representative of two independent experiments.

Motivated by this analysis, we tested whether HDAC active site inhibitors could block CoREST degradation by KBTBD4 MB mutants (**Fig. 6b**). Treatment with SAHA and CI-994, a 2- amino-benzamide-derived inhibitor, could rescue CoREST degradation induced by nearly all MB ligase mutants. The notable exception was the failure of SAHA to block the activity of KBTBD4- PR, highlighting the potential plasticity of the E3-substrate interaction. Nonetheless, the activity of KBTBD4-PR could be inhibited by CI-994, likely due to its larger size and/or its slower off-rate^33^. We further confirmed that CI-994 could block association of KBTBD4-P and KBTBD4-PR with HDAC1 in cells as well as in a purified reconstituted system (**Fig. 6c, d**). We next tested a more recently developed selective HDAC1/2-inhibitor, RBC1HI^34^. Consistent with its even larger size, RBC1HI was effective at blocking CoREST degradation (**Fig. 6b**) but showed reduced activity in our TR-FRET assay, especially against the more potent KBTBD4-PR mutant (**Fig. 6d**). Although seemingly incongruous, these findings are consistent with past studies showing that selective HDAC1/2 inhibitors possess slow-binding kinetics and often require disassociation of the corepressor complex to engage HDAC1/2 with high affinity^24,35^, which may not occur on the shorter time scale of the in vitro experiments. Together, these findings support the idea that selective HDAC1/2 inhibitors can disengage mutant KBTBD4 from HDAC1/2 to stabilize the associated corepressor complexes.

Our results suggest that HDAC1/2-selective inhibitors, such as RBC1HI^34^, could selectively block the proliferation of KBTBD4-mutant MB tumor cells. To explore this concept, we evaluated the sensitivity of MB PDX models harboring KBTBD4-PR to RBC1HI treatment. Ex vivo treatment of dissociated, resected tumor cells from mice transplanted with either WT (MED411, *n* = 2 mice; RCMB28, *n* = 3 mice) or KBTBD4-PR mutant (ICB1572, *n* = 5 mice) PDX cells showed that the mutant displayed significantly heightened sensitivity to RBC1HI (**Fig. 6e**). Moreover, ex vivo treatment of ICB1572 cells with MLN4924 and RBC1HI, separately, led to increased levels of CoREST and LSD1, consistent with KBTBD4 inhibition (**Fig. 6f**). Altogether, these results demonstrate the potential promise of selective HDAC1/2 inhibition as a strategy to block the growth of KBTBD4-mutant MB cells.

## Discussion

Here, we elucidate the molecular mechanism by which hotspot cancer mutations in the E3 ligase KBTBD4 can reprogram its protein-protein interactions to promote aberrant degradation of HDAC1/2 corepressor complexes in MB, implicating HDAC1/2 as the relevant target of mutant KBTBD4 for the first time. Using deep mutational scanning, we unveil the mutational landscape and molecular rules that control this neomorphic activity, highlighting how insertion mutations fundamentally differ from point substitutions in their preferences, effects, and cooperativity, albeit within this specific E3 ligase context. Moreover, these findings underscore how insertions at a protein surface, in comparison to point substitutions, can be particularly effective at promoting neo-PPIs.^28^ Leveraging these data with cryo-EM, we reveal the mechanistic basis by which MB mutations reconfigure the 2b-2c loop of KBTBD4 to extend into the HDAC1 active site in a shape-complementary fashion. Remarkably, these additional contacts made by the cancer mutations precisely mimic the effects of UM171 in gluing the suboptimal KBTBD4-HDAC1 interface, showcasing how chemical and human genetic perturbations can act as molecular facsimiles. Lastly, understanding of this molecular interface establishes the rationale for employing HDAC1/2 inhibitors to block the activity of KBTBD4 MB mutants and the growth of *KBTBD4*-mutant PDX MB models. Further studies will be required to fully investigate the therapeutic potential of this approach and the downstream role of HDAC1/2 corepressor degradation in MB tumorigenesis.

We have recently shown that most molecular glue compounds operate by potentiating weak, intrinsic interactions between two target proteins^36^. In agreement, WT KBTBD4 shows low basal affinity towards LHC (**Fig. 1e**). By complementing the KBTBD4-HDAC1 interface, both UM171 and the MB cancer mutations can increase the protein binding affinity > 5-10-fold. Similar to small molecule molecular glues, cancer mutations, therefore, can also exploit and convert non-productive basal protein-protein interactions to confer neomorphic functions. This mechanistic convergence of a molecular glue and human genetic mutations demonstrates how these perturbations can operate by similar molecular principles, and we anticipate that future instances of this chemical genetic paradigm may be uncovered and exploited for therapeutic applications. In conclusion, our study defines the mechanistic basis of E3 ligase gain-of-function cancer mutations for the first time and raises the prospect for how massively parallel genetic methods may eventually enable *de novo* molecular glue discovery and design by identifying “glueable” protein sites.

## Acknowledgements

The authors would like to thank J. Nelson at the Bauer Core Facility of Harvard University for assistance with FACS, J.D. Quispe and S. Dickinson at the Arnold and Mabel Beckman Cryo-EM Center of the University of Washington, R. Yan, X. Zhao, J. Jung, and Z. Yu at the Cryo-EM Facility on the Janelia Research Campus of the Howard Hughes Medical Institute, and T. Humphreys and M. Campbell at Fred Hutch EM & cryo-EM Core for their assistance in electron microscopy data acquisition, as well as D. Asarnow from the Veesler laboratory at the University of Washington for his technical insights and suggestions, and members of the Zheng and Liau laboratories, especially D.V. Rusnac, S. Zhang, H. Shi, E. Garcia, J. Woods, S. Shen, and N. Lue, for their discussion and inputs. We also acknowledge the Center for Proteomics and Metabolomics and the Animal Resources Center at St. Jude for their support. N.C.P is supported by the National Science Foundation (DGE1745303). P.A.C. is supported by the National Institute of General Medical Sciences (R35GM149229) and the Leukemia and Lymphoma Society. P.A.N. is supported by the American Lebanese Syrian Associated Charities (St. Jude), The Brain Tumor Charity (Quest for Cures), The Mark Foundation (Emerging Leader Award), Alex’s Lemonade Stand Foundation (Crazy 8 Initiative), and the National Cancer Institute (P01CA096832-16A1; 1R01CA270785-01A1). N.Z. is supported by the Howard Hughes Medical Institute. B.L. is supported by the Ono Pharma Foundation, the Camille and Henry Dreyfus Foundation (Teacher-Scholar Award), the Blavatnik Accelerator Fund (Harvard University), the National Institute of General Medical Sciences (1DP2GM137494), and the National Cancer Institute (1R01CA274437).

## Author contributions

B.L., V.D., and N.Z. conceived the project with inputs from X.X., O.Z., and M.Y.. O.Z. performed cellular experiments and deep mutational scanning, and purified KBTBD4 for biochemical studies. M.Y. performed cellular experiments and flow cytometry. C.L. conducted computational analysis for deep mutational scanning. O.Z. and E.N. purified LHC for biochemical and structural studies and H.J. conducted in vitro ubiquitination experiments, all with input from P.A.C.. O.Z. and N.C.P. conducted TR-FRET experiments with input from R.M.. S.H. synthesized UM171 and RBC1HI. H.S.K. assisted with cellular experiments and deep mutational scanning. L.P. performed PDX transplantation and ex vivo cell assays with support from J.H., M.B., and P.A.N.. Y.L. analyzed MB proteomics data generated with support from J.H., H.L., and P.A.N.. H.M. and X.X. purified KBTBD4 mutants for structural studies and CUL3-RBX1 for in vitro ubiquitination assays. X.X. performed cryo-EM grid preparation, specimen screening, data collection and processing. X.X., N.Z., and B.L. analyzed the structures. N.Z. and B.L. held overall responsibility for the study.

## Supplementary Materials

Materials and Methods

Extended Data Figures 1 to 7

Extended Data Table 1

Supplementary Note 1 | Synthetic procedures

Supplementary Note 2 | NMR Spectra

Supplementary Table 1 | Primers used for DMS library cloning

Supplementary Table 2 | Primers used for DMS plasmid construction

Supplementary Table 3 | Primers used for DMS sequencing

Supplementary Data 1 | KBTBD4 MUT vs WT PDX proteomics data

Supplementary Data 2 | DMS library sequences

Supplementary Data 3 | DMS enrichment scores

## Materials and Methods

### MB patient-derived xenograft (PDX) lines

Ex vivo drug screening and TMT-proteomic study on patient-derived xenografts (PDX) were performed at St. Jude Children’s Research Hospital (SJCRH). NSG mice (NOD.Cg- *Prkdc^scid^Il2rg^tm1Wjl^/SzJ* (The Jackson Laboratory, JAX#005557) were used as hosts for PDX studies. Female NSG mice at least 8 weeks of age were anesthetized in a surgical suite, and dissociated PDX cells were implanted in the cerebellum to amplify tumor material for down- stream analyses. Mice were observed daily and euthanized at the onset of signs of sickness, including lethargy and neurological abnormalities. All clinical signs at the time of euthanasia did not exceed humane endpoint as determined by SJCRH Institutional Animal Care and Use Committee (IACUC protocol#589-100536). RCMB51, RCMB52, and RCMB28 were originated and shared by Robert J. Wechsler-Reya, Ph.D., Columbia University (previously Stanford). ICB1299 and ICB1572 were originated and shared by Xiao-Nan Li, M.D., Ph.D., Northwestern University Feinberg School of Medicine (previously Baylor University). MED411, MED211, and MED2312 were purchased from Brain Tumor Research Laboratory, Seattle Children’s Hospital (previously Fred Hutchinson)^37^. Low passage PDXs (<10) were dissected and then flash frozen for proteomics or dissociated for *ex vivo* drug sensitivity screening.

### Sample Processing of mouse PDX tissues for TMT-MS

Frozen tissues (20-30 mg) from each mouse PDX tumor were added to 200 mL freshly prepared 8M Urea lysis buffer (containing 12 g Urea, 10X HEPES in 25 mL Millipore ultrapure water), homogenized with glass beads in Bullet Blender Tissue Homogenizer (Next Advance Inc.) for 5 min, followed by a 2 min centrifugation at 2000 rpm. Subsequently, 1% sodium deoxycholate was immediately added to the lysed tissues and vortexed for 2 min, followed by centrifugation at 1000 rpm. The resulting supernatants were collected and stored at –80 °C. For quality control and quantification, 2 mL lysates from each sample were electrophoresed on 4-12% NuPAGE gels (Invitrogen)^38^.

### Protein Digestion and Tandem-Mass-Tag (TMT) Labeling

The analysis was performed with a previously optimized protocol^38,39^. For whole proteome profiling, quantified protein samples (300 µg in the lysis buffer with 8 M urea) for each TMT channel were proteolyzed with Lys-C (Wako, 1:100 w/w) at 21 °C for 2 h, diluted by 4-fold to reduce urea to 2 M for the addition of trypsin (Promega, 1:50 w/w) to continue the digestion at 21 °C overnight. The insoluble debris was kept in the lysates for the recovery of insoluble proteins. The digestion was terminated by the addition of 1% trifluoroacetic acid. After centrifugation, the supernatant was desalted with the Sep-Pak C18 cartridge (Waters), and then dried by Speedvac (Thermo Fisher). Each sample was resuspended in 50 mM HEPES (pH 8.5) for TMT labeling and then mixed equally, followed by desalting for the subsequent fractionation. For the whole proteome analysis alone, 0.1 mg protein per sample was used.

### Extensive Two-Dimensional Liquid Chromatography-Tandem Mass Spectrometry (LC/LC-MS/MS)

The TMT labeled samples were fractionated by offline basic pH reverse phase LC, and each of these fractions was analyzed by the acidic pH reverse phase LC-MS/MS^40,41^. We performed a 160 min offline LC run at a flow rate of 400 µL/min on an XBridge C18 column (3.5 μm particle size, 4.6 mm x 25 cm, Waters; buffer A: 10 mM ammonium formate, pH 8.0; buffer B: 95% acetonitrile, 10 mM ammonium formate, pH 8.0)^38^. A total of 80 two-minute fractions were collected. Every 41^st^ fraction was concatenated into 40 pooled fractions, which were subsequently used for whole proteome TMT analysis.

In the acidic pH LC-MS/MS analysis, each fraction from basic pH LC was dried by a Speedvac and was run sequentially on a column (75 µm x 35 cm for the whole proteome, 50 µm x 30 cm for whole proteome, 1.9 µm C18 resin from Dr. Maisch GmbH, 65 °C to reduce backpressure) interfaced with a Fusion MS (Thermo Fisher) for the whole proteome where peptides were eluted by a 90 min gradient (buffer A: 0.2% formic acid, 5% DMSO; buffer B: buffer A plus 65% acetonitrile). MS settings included the MS1 scan (410-1600 m/z, 60,000 resolution, 1 x 10^6^ AGC and 50 ms maximal ion time) and 20 data-dependent MS2 scans (fixed first mass of 120 m/z, 60,000 resolution, 1 x 10^5^ AGC, 200 ms maximal ion time, HCD, 36% normalized collision energy, 1.0 m/z isolation window with 0.2 m/z offset, and 20 s dynamic exclusion). MS settings included the MS1 scan (450-1600 m/z, 60,000 resolution, 1 x 10^6^ AGC and 50 ms maximal ion time) and 20 data-dependent MS2 scans (fixed first mass of 120 m/z, 60,000 resolution, 1 x 10^5^ AGC, 110 ms maximal ion time, HCD, 36% normalized collision energy, 1.0 m/z isolation window with 0.2 m/z offset, and 10 s dynamic exclusion).

### Proteins Identification and Quantification with JUMP Software

The computational processing of identification and quantification was performed with the JUMP search engine^40^. All original target protein sequences were reversed to generate a decoy database that was concatenated to the target database. Putative PSMs were filtered by mass accuracy and then grouped by precursor ion charge state and filtered by JUMP-based matching scores (Jscore and ΔJn) to reduce FDR below 1% for proteins during the whole proteome analysis. If one peptide could be generated from multiple homologous proteins, based on the rule of parsimony, the peptide was assigned to the canonical protein form in the manually curated SwissProt database. If no canonical form was defined, the peptide was assigned to the protein with the highest PSM number. The analysis was performed in the following steps, as previously reported with modifications^42^: (i) extracting TMT reporter ion intensities of each PSM; (ii) correcting the raw intensities based on the isotopic distribution of each labeling reagent (e.g. TMT126 generates 91.8%, 7.9% and 0.3% of 126, 127, 128 m/z ions, respectively); (iii) excluding PSMs of very low intensities (e.g. minimum intensity of 1,000 and median intensity of 5,000); (iv) removing sample loading bias by normalization with the trimmed median intensity of all PSMs; (v) calculating the mean-centered intensities across samples (e.g. relative intensities between each sample and the mean), (vi) summarizing protein or phosphopeptide relative intensities by averaging related PSMs; (vii) finally deriving protein or phosphopeptide absolute intensities by multiplying the relative intensities by the grand-mean of three most highly abundant PSMs. In addition, we also performed y1-ion based correction of TMT data. See Supplementary Data 1.

### Differential Expressed Analysis of Proteins

Differentially expressed proteins (DEPs) were identified using the limma package (version 3.54.2)^43^. Low expressions were defined as the lower 25^th^ percentile of the means of the protein expression, and proteins with a prevalence of low expression in more than 70% of the samples were filtered out. As a result, 7731 out of 11428 proteins were retained for further analysis. Criteria for differential expression included a p-value < 0.01 and a fold change > 1.5. Protein-protein interaction networks were constructed using STRINGdb (version 12)^44^, with a confidence threshold >0.7. Interaction data were sourced from text mining, experiments, and existing databases.

### Cell culture

HEK293T cells were a gift from B.E. Bernstein (Massachusetts General Hospital). K562 cells were obtained from ATCC. All mammalian cell lines were cultured in a humidified 5% CO_2_ incubator at 37 °C and routinely tested for mycoplasma (Sigma-Aldrich). HEK293F cells were obtained from Thermo Fisher. RPMI1640 and DMEM were supplemented with 100 U mL^−1^ penicillin and 100 µg mL^−1^ streptomycin (Gibco) and FBS (Peak Serum). K562 cells were cultured in RPMI1640 (Gibco) supplemented with 10% FBS. HEK293T cells were cultured in DMEM (Gibco) supplemented with 10% FBS. HEK293F cells were cultured in Freestyle™ 293 Expression Medium (Thermo Fisher) shaking at 125 rpm. *Spodoptera frugiperda* (Sf9) insect cells (Expression Systems, 94-001F) were cultured in ESF921 media (Expression Systems) in a non-humidified and non-CO_2_ incubator at 27 °C shaking at 140 rpm. High Five cells were purchased from Thermo Fisher (B85502), with Grace insect medium (Thermo Fisher, 11595030) supplemented with 10% FBS (Cytiva) and 1% Penicillin-Streptomycin (Gibco), cultured at 26 °C.

### Lentiviral production

For lentivirus production, transfer plasmids were co-transfected with GAG/POL and VSVG plasmids into 293T cells using Lipofectamine™ 3000 (Thermo Fisher Scientific) according to the manufacturer’s protocol. Media was exchanged after 6 h and the viral supernatant was collected 52 h after transfection and sterile-filtered (0.45 µm). K562 cells were transduced by spinfection at 1,800 × *g* for 1.5 h at 37 °C with 8 µg mL^-1^ polybrene (Santa Cruz Biotechnology).

### Plasmid Construction

Plasmids were cloned by Gibson Assembly using NEBuilder HiFi (New England Biolabs). Cloning strains used were NEB Stable (lentiviral) (New England Biolabs). Final constructs were validated by Sanger sequencing (Azenta/Genewiz).

All KBTBD4 expression plasmids encoded isoform 1 (human, residues 1-518) but longer isoform 2 (residues 1-534) numbering was used. CoREST expression plasmids encoded isoform 1 (human) full length (residues 1-482). ORFs of human KBTBD4 and CoREST (mammalian expression) were amplified from ORFs obtained from Horizon Discovery.

For transfection constructs, CoREST-FLAG and HA-KBTBD4 (WT or mutant) constructs were cloned into pcDNA3. For KBTBD4 overexpression constructs, KBTBD4 coding sequences were cloned into pSMAL mCherry, which was generated from pSMAL through introduction of an mCherry ORF into pSMAL (a gift from J. E. Dick, University of Toronto). For bacmid expression, KBTBD4 was cloned into pFastbac, a gift from T. Cech.

### CRISPR/Cas9-mediated genome editing

mEGFP followed by a ‘GGGSGGGS’ linker was knocked into the C-terminus of CoREST (i.e., *RCOR1*) in K562 cells. sgRNA (sgRNA: TTCAAAGCCACCAGTTTCTC) targeting the C-terminus of CoRESTwas cloned into a Cas9 plasmid, PX459^45^, and electroporated according to the manufacturer’s protocol (Neon™ Transfection System, Thermo Fisher Scientific) with a repair vector containing the mEGFP CDS and linker flanked by 750 bp of genomic homology sequences to either side of the CoREST C-terminus. Briefly, 2 x 10^5^ cells were washed twice with PBS and resuspended in buffer R. PX459 (0.5 µg) and the repair vector (0.5 µg) were added to the cell suspension, and electroporated at 1350 V with 10 ms pulse width for 4 pulses using the Neon^TM^ Transfection System 10 µL kit. After electroporation, cells were immediately transferred to prewarmed media. To generate single-cell clones, cells were gated to sort for the top 0.2% GFP^+^ and single-cell sorted on a MoFlo Astrios EQ Cell Sorter (Beckman Coulter), expanded, and validated by western blot and Sanger sequencing.

### Generation of knockout K562s

KBTBD4 KO CoREST-GFP K562 clones were generated by using the Alt-R™ CRISPR-Cas9 System (IDT) to deliver ribonucleoprotein complexes containing the KBTBD4 KO guide (KBTBD4: GATATCTGTGAGTAAGCGGT) using the Neon™ Transfection System (Thermo Fisher Scientific) according to the manufacturer’s protocol. Transfected cells recovered for 72 h before sorting for single cell clones on a MoFlo Astrios Cell Sorter (Beckman Coulter). Single cell clones were validated by genotyping and immunoblotting.

### Degradation assay of KBTBD4 mutants

K562 KBTBD4-null CoREST-GFP cells were generated as described above. KBTBD4 overexpression constructs were cloned into pSMAL mCherry and point mutations were introduced into coding regions using standard PCR-based site-directed mutagenesis techniques. Lentiviral particles carrying the overexpression constructs were produced and used to transduce K562 KBTBD4-null CoREST-GFP cells as described above. 48 h after transduction, cells were treated with 1 µM UM171, 1 µM MLN4924, 10 µM HDACi (SAHA, CI-994, or RBC1HI), or 0.1% vehicle for 24 h. GFP^+^% was measured for mCherry^+^ cells in each condition (**Extended Fig. 2c**).

### Protein expression and purifications

Human recombinant KBTBD4 for biochemical and biophysical analyses was purified from Sf9 insect cells. cDNAs for human KBTBD4 was cloned into the pFastBac donor vector and the recombinant baculovirus was constructed using the Bac-to-Bac protocol and reagents (Thermo Fisher Scientific). KBTBD4 medulloblastoma mutations were introduced into coding regions using standard PCR-based site-directed mutagenesis techniques. All KBTBD4 constructs were tagged on the N-terminus with 6×His cleavable by TEV protease. These plasmids were used to prepare separate baculoviruses according to standard protocols (Bac-to-Bac Baculovirus Expression System, Thermo Fisher). Detection of gp64 was used to determine baculovirus titer (Expression Systems). For expression, SF9 cells were grown to a density of 1-2 × 10^6^ cells/mL and infected with KBTBD4 baculovirus. The cells were incubated for 72 h (27 °C, 120 × *g*), harvested and then frozen with liquid nitrogen for future purification. Cells were resuspended in lysis buffer (50 mM Tris-HCl, pH 8.0 cold, 500 mM NaCl, 1 mM TCEP, 10% glycerol, 15 mM imidazole) supplemented with 1% NP-40, 1 mM PMSF, and Roche Complete Protease Inhibitor and sonicated. Lysate was clarified by centrifugation at 100,000 × *g* for 30 min and incubated with His60 Ni Superflow affinity resin (Takara). Resin was washed with lysis buffer containing a stepwise gradient of 15–50 mM imidazole, followed by elution using lysis buffer with 250 mM imidazole. Eluate was exchanged into storage buffer (50 mM Tris-HCl, pH 8.0 cold, 150 mM NaCl, 1 mM TCEP, 10% glycerol) using an Econo-Pac 10DG desalting column (Bio-Rad) and further purified by size exclusion chromatography using a Superdex 200 10/300 GL column (GE Healthcare). The purity of the recombinant protein was verified by SDS-PAGE and fractions with 90–95% purity were pooled and stored at –80 °C.

Recombinant human KBTBD4 used in cryo-EM structure determination was purified from *Trichoplusia ni* High Five insect cells. cDNAs for human KBTBD4 was cloned into the pFastBac donor vector and the recombinant baculovirus was constructed using the Bac-to-Bac protocol and reagents (Thermo Fisher Scientific). KBTBD4 constructs were tagged on the N-terminus with 10xHis and MBP tag cleavable by TEV protease. These plasmids were used to prepare separate baculoviruses according to standard protocols (Bac-to-Bac Baculovirus Expression System, Thermo Fisher). For expression, the monolayer High Five cells were grown to about 80% confluency and infected with KBTBD4 baculovirus. The cells were incubated for 72 h (26 °C), harvested, and then frozen with liquid nitrogen for future purification. Cells were resuspended in lysis buffer (50 mM Tris-HCl, pH 8.0 cold, 150 mM NaCl, 1 mM TCEP, 20 mM imidazole) supplemented with 1 mM PMSF, 10 µM Leupeptin, 0.5 µM Aproptinin and 1 µM Pepstatin A and sonicated. Lysate was clarified by centrifugation at 100,000 × *g* for 30 min and incubated with amylose affinity resin (New England BioLabs). Resin was washed with lysis buffer, followed by elution using lysis buffer with 10 mM maltose. Eluate was cut with Tobacco Etch Virus protease overnight, followed by the prepacked anion exchange column (GE Healthcare) to get rid of the protease and further purified by size exclusion chromatography using a Superdex 200 10/300 GL column (GE Healthcare). The purity of the recombinant protein was verified by SDS-PAGE and fractions with 90–95% purity were pooled and stored at –80°C.

Recombinant LSD1-CoREST-HDAC complex was comprised of full length LSD1 (UniProt ID: O60341) or LSD1 (Δ77-86), full length HDAC1 (UniProt ID: Q13547) and N-terminally truncated CoREST (aa 84-482) (UniProt ID: Q9UKL0) or N-terminal Cys CoREST^22^. The pcDNA3 vector was used to create plasmids encoding the different proteins. The CoREST constructs contained an N-terminal (His)10(Flag)3 tag followed by a TEV protease cleavage site. The constructs for ternary complex were co-transfected into suspension-grow HEK293F cells (ThermoFisher Scientific) with polyethylenimine (PEI) (Sigma) and harvested after 48 h. Cells were resuspended in lysis buffer (50 mM HEPES, pH7.5, 100 mM KCl, 5% glycerol, 0.3% Triton X-100, 1X Roche EDTA-free Complete Protease Inhibitor cocktail) and sonicated. Lysate was clarified by centrifugation at 12,000 × *g* for 30 min and incubated with Anti-Flag M2 affinity gel (Sigma). The affinity gel was washed twice with lysis buffer and twice with SEC buffer (50 mM HEPES, pH 7.5, 50 mM KCl, 0.5 mM TCEP) followed by the incubation with TEV protease overnight at 4 °C. The complex was further purified by size exclusion chromatography using a Superose 6 10/300 column (GE Healthcare). The purity of the complex was verified by SDS-PAGE and fractions with 90–95% purity were pooled and supplemented with 5% glycerol and stored at –80 °C.

### Fluorescein labeling of LHC

The fluorescein labeling of the LSD1-CoREST-HDAC1 complex was purified as described above. A Cys point mutagenesis has been conducted next to the TEV protease cleavage site of N- terminally truncated CoREST for the ligation reaction with NHS- fluorescein ^46^. A 2 mM NHS-fluorescein was incubated with 500 mM mercaptoethanesulfonate (MESNA) in the reaction buffer (100 mM HEPES, pH 7.5, 50 mM KCl, 1 mM TCEP) for 4 h at room temperature in the dark for transesterification. The LSD1-CoREST-HDAC1 complex purified by FLAG M2 affinity gel was washed with reaction buffer and incubated with TEV protease for 5 h at 4 °C. The complex was then mixed with 500 µL of the fluorescein/MESNA solution to make a final concentration of 0.5 mM fluorescein and 125 mM MESNA. The mixture was incubated for 48 h at 4 °C in the dark. The complex was desalted by a Zeba spin desalting column (7 kDa MWCO) and further purified by size exclusion chromatography using a Superose 6 10/300 column (GE Healthcare). Fluorescein-labeling efficiency was analyzed by SDS-PAGE and fluorescence gel imaging (Amersham Typhoon FLA 9500, Cytiva). The purity of the complex was verified by SDS-PAGE and fractions with 90–95% purity were pooled and supplemented with 5% glycerol and stored at –80 °C.

### TR-FRET measurements

Unless otherwise noted, experiments were performed in white, 384-well microtiter plates (Corning 3572) in 30 μL assay volume, or white, 384-well low-volume microtiter plates (PerkinElmer 6008280). TR-FRET measurements were acquired on a Tecan SPARK plate reader with SPARKCONTROL software version V2.1 (Tecan Group Ltd.), with the following settings: 340/50 nm excitation, 490/10 nm (Tb), and 520/10 nm (FITC, AF488) emission, 100 μs delay, 400 μs integration. The 490/10 nm and 520/10 nm emission channels were acquired with a 50% mirror and a dichroic 510 mirror, respectively, using independently optimized detector gain settings unless specified otherwise. The TR-FRET ratio was taken as the 520/490 nm intensity ratio on a per-well basis.

### Ternary complex measurements by TR-FRET

#### Titration of Fluorescein-labeled LSD1-CoREST-HDAC complex

Recombinant WT (or mutant) 6×His-KBTBD4 (20 nM, 2×) and CoraFluor-1-labeled anti-6×His IgG (10 nM, 2×)^25^ were diluted into LHC buffer, with or without 10 μM UM171, and 5 μL added to wells of a white, 384-well low volume microtiter plate (PerkinElmer 6008280). Serial dilutions of fluorescein-labeled LSD1-CoREST-HDAC complex (1:2 titration, 10-point, c_max_ = 1,000 nM, 2×) were prepared in ligand buffer and 5 μL added to wells of the same plate (final volume 10 μL, final 6×His-KBTBD4 concentration 10 nM, final CoraFluor-1-labeled anti-6×His IgG concentration 5 nM, fluorescein-labeled LSD1-CoREST-HDAC complex c_max_ 500 nM). The plate was allowed to equilibrate for 1 h at room temperature before TR-FRET measurements were taken. Data were background-corrected from wells containing no 6×His-KBTBD4. Prism 9 was used to fit the data to a four-parameter dose-response curve.

#### Titration of InsP_6_

Recombinant WT or mutant (P and PR) 6×His-KBTBD4 (40 nM), fluorescein-labeled LSD1- CoREST-HDAC complex (40 nM), and CoraFluor-1-labeled anti-6×His IgG (20 nM)^25^ were diluted into were diluted into a one-to-one mixture of ligand buffer (50 mM Tris-HCl, pH 8.0, 150 mM NaCl, 1 mM TCEP, 10% glycerol) and LHC buffer (20 mM HEPES, pH 7.5, 1 mM TCEP, 2 mg/mL BSA, 0.1% Tween-20) and 10 μL added to wells of a white, 384-well low volume microtiter plate (PerkinElmer 6008280). InsP_6_ was added in serial dilution (1:10 titration, 6-point, c_max_ = 100 μM) using a D300 digital dispenser (Hewlett-Packard), and allowed to equilibrate for 1 h at room temperature before TR-FRET measurements were taken. Data were background-corrected from wells containing no InsP_6_. Prism 9 was used to fit the data to a four-parameter dose-response curve.

#### Incubation with HDAC inhibitor

Fluorescein-labeled LSD1-CoREST-HDAC complex (100 nM), and CoraFluor-1-labeled anti- 6×His IgG (20 nM)^25^ were diluted into were diluted into a one-to-one mixture of ligand buffer (50 mM Tris-HCl, pH 8.0, 150 mM NaCl, 1 mM TCEP, 10% glycerol) and LHC buffer (20 mM HEPES, pH 7.5, 1 mM TCEP, 2 mg/mL BSA, 0.1% Tween-20, 100 µM InsP_6_) and 10 μL added to wells of a white, 384-well low volume microtiter plate (PerkinElmer 6008280). HDACi (SAHA, CI994, RBC1HI) (10 µM) or vehicle (DMSO) was added using a D300 digital dispenser (Hewlett-Packard), and allowed to equilibrate for 1 h at room temperature. Recombinant WT or mutant (P or PR) 6×His-KBTBD4 (100 nM) was then added using a D300 digital dispenser (Hewlett-Packard), and allowed to equilibrate for 1 h at room temperature before TR-FRET measurements were taken. Data were background-corrected from wells containing no 6×His-KBTBD4. Prism 9 was used plot the data.

### *In vitro* ubiquitination assay

The ubiquitination assays were set up similarly as previously reported^47^. Reactions were performed at 37°C in a total volume of 20 µL. The reaction mixtures contained 5 mM ATP, 100 μM WT ubiquitin, 100 nM E1 protein, 2 μM E2 protein, 0.5 μM neddylated RBX1-CUL3, 0.5 µM WT or PR KBTBD4 (unless otherwise indicated) with 25 mM Tris-HCl (pH 7.5), 20 mM NaCl, 10 µM InsP_6_, and 2.5 mM MgCl2 as reaction buffer. Substrate LHC at 0.5 µM was preincubated with everything except E1 in the reaction mixture at 37°C for 5 min prior to adding E1 to initiate the reaction. Reactions were quenched at the indicated time points by adding SDS loading buffer containing reducing agent β-mercaptoethanol. The reaction samples were resolved on SDS-PAGE gels and analyzed by Colloidal Blue staining, western blots, or Typhoon fluorescent imaging.

### Deep mutational scan

The library of KBTBD4 mutants in the 7aa region between Gly307 and Arg313 was designed to comprise all possible: i) 1 aa deletions ii) 1 aa substitutions iii) 2 aa substitutions of adjacent residues iv) 1 aa insertions v) 2 aa insertions (2,680) vi) 3aa insertions of GGG or GSG vii) 100 randomly scrambled WT sequences and the 2 remaining MB indels (PR311delinsPPHV, IPR310delinsTTYML). The 5’ and 3’ homology arms were added as well as forward and reverse barcodes for the different sub pools of mutations for downstream cloning. The final library was ordered from Twist Biosciences as a pooled oligo library with final lengths of single stranded oligos ranging from 101-113nt (Supplementary Data 2). The Twist pool was resuspended in TE to the concentration of 1 ng/uL and the sub pools were separated by PCR amplification of 22 cycles, using lsPCR1 primers listed in Supplementary Table 1 and using 1ng of the Twist pool as template in each reaction. Each sub pool was further amplified with lsPCR2 primers in Supplementary Table 1 by PCR amplification of 10 cycles and the library pools were gel purified (Zymo Gel DNA Recovery Kit).

The mutational library was cloned into pSMAL mCherry using Gibson assembly. The backbone for the Gibson assembly was prepared by introducing a BamHI restriction site in place of residues Gly307 and Arg313 using primers in Supplementary Table 2. The backbone was digested with BamHI (NEB) and subsequently treated with Antarctic phosphatase (NEB) and the correct linearized backbone was isolated by gel electrophoresis and purified using Gel DNA Recovery Kit (Zymo). 190 ng of linearized vector and 13.15 ng of each sub pool were used for each Gibson reaction of 80 µL using HIFI DNA Assembly Master Mix (NEB). The Gibson reaction was incubated for 1 hr at 50°C and DNA was isolated by isopropanol precipitation and transformed into Lucigen Endura Competent Cells according to manufacturer’s protocol. Cells recovered in Lucigen Endura Recovery Media for 1 hr at 30°C and later plated and grown overnight at 30°C. Colonies were harvested and the plasmid library was extracted using QIAGEN Plasmid Maxi Kit. Purified sub pools were then combined for the final library and sequence verified on an Illumina MiSeq as previously described.

Lentivirus was produced and titered by measuring cell counts after transduction and mCherry selection. K562 KBTBD4-null CoREST-GFP knock-in cells were transduced with library lentivirus at a multiplicity of infection <0.3 and at day 3 post transduction, cells were sorted on a MoFlo Astrios Cell Sorter (Beckman Coulter), collecting top 10% GFP^-^ and mCherry^+^, GFP^+^ and mCherry^+^, and mCherry^+^ (GPF^+/-^) cells. Genomic DNA was isolated using the QIAamp DNA Blood Mini kit or QIAamp UCP DNA Micro kit, and mutation sequences were amplified using barcoded primers listed in Supplementary Table 3, purified by gel extraction and sequenced on an Illumina MiSeq as previously described. Sorting was performed in 3 reps and at all steps, >150X coverage of the library was maintained.

Data analysis was performed using Python (v.3.9.12) with Biopython (v.1.78), Pandas (v.1.5.1) and NumPy (v.1.23.4). Briefly, raw reads matching sequences in the mutational library from unsorted as well as sorted (GFP^+^ and GFP^-^) cells were counted. Counts were then processed by converting them to reads per million, adding a pseudocount of 1 and transforming them by log_2_. Enrichment of each variant in GFP^+^ and GFP^-^ populations were quantified by subtracting the GFP^+^ and GFP^-^ log_2_-transformed counts, respectively, by corresponding log_2_-transformed counts for unsorted cells and averaged across replicates (Supplementary Data 3). Heatmaps were generated using matplotlib (v3.7.1).

### Analysis of sequence motifs

Position probability matrices of the GFP^+^ and unsorted population were constructed for each mutually exclusive category (single substitution, single insertion, double substitution, and double insertion) by normalizing raw counts by the total read counts of each corresponding category, averaging across replicates. and tallying the probability of every amino acid at each position. The information content *IC* of each position *N* were calculated according to Kullback-Leibler divergence, which is as follows:

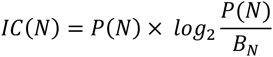

Where, P(N) is the position probability matrices of GFP^+^ population for each mutational category and the position probability matrix of the unsorted population was used as background frequencies B_N_. Logos were generated using Logomaker (v0.8)^48^.

### Immunoblotting

Cells were lysed on ice in RIPA buffer (Boston BioProducts) with 1X Halt Protease Inhibitor Cocktail (Thermo Fisher Scientific) and 5 mM EDTA (Thermo Fisher Scientific). Lysate was clarified by centrifugation and total protein concentration was measured with the BCA Protein Assay (Thermo Fisher Scientific). Samples were electrophoresed and transferred to a 0.45-μm nitrocellulose membrane (Bio-Rad). Membranes were blocked with tris-buffered saline Tween (TBST) with 5% Blotting-Grade Blocker (Bio-Rad) and incubated with primary antibody: KBTBD4 (Novus Biologicals, catalog no. NBP1-88587, 1:1,000), HDAC1 (Cell Signaling Technology, catalog no. 34589, D5C6U, 1:1,000), FLAG (Sigma-Aldrich, catalog no. F1804, M2, 1:2,000), His-tag (Cell Signaling Technology, catalog no. 2365, 1:1,000), HA-tag (Cell Signaling Technology, catalog no. 3724, C29F4, 1:1,000), GAPDH (Santa Cruz Biotechnology, catalog no. sc-47724, 0411, 1:10,000). Membranes were washed 3X with TBST and incubated with secondary antibody: anti-rabbit IgG HRP conjugate (Promega, catalog no. W4011, 1:20,000), anti-mouse IgG HRP conjugate (Promega, catalog no. W4021, 1:40,000). Unless otherwise stated, following 3X washes with TBST, immunoblots were visualized using SuperSignal West Pico PLUS or SuperSignal West Femto chemiluminescent substrates (Thermo Fisher Scientific).

### Co-immunoprecipitation

HEK293T cells were transfected with 2 µg pcDNA3 HA-KBTBD4 plasmid (mutant or WT) and with or without 3 µg pcDNA3 CoREST-FLAG (full-length or truncated) using PEI MAX transfection reagent (Polysciences) according to the manufacturer’s protocol. 48 h after transfection, cells were treated with 1 µM MLN4924 for 3 h then 1 µM UM171, 10 µM CI-994, or vehicle for 1 h. Cells were washed twice with cold PBS and flash frozen.

Cells were thawed, lysed on ice in lysis buffer (25 mM Tris-HCl pH 7.5, 150 mM NaCl, 1% NP-40 alternative) supplemented with cOmplete™, EDTA-free Protease Inhibitor Cocktail (Sigma-Aldrich), and the lysates were cleared. The protein concentration was quantified as above and diluted to 1 mg ml^−1^ in lysis buffer with 1 µM UM171 or DMSO. Supernatants were immunoprecipitated overnight at 4 °C with 25 µL Pierce anti-HA magnetic beads (Thermo Fisher Scientific). Beads were washed six times with lysis buffer, eluted in SDS-PAGE loading buffer, and carried forward to immunoblotting as described above.

### Ex vivo drug sensitivity screening in MB PDX cells

MB patient derived xenografts harboring wild type KBTBD4 (RCMB28 *n* = 3, MED411 *n* = 2) or KBTBD4-PR mutant (ICB1572 *n* = 5) were used to assess sensitivity to the HDAC1/2 inhibitor RBC1HI. Briefly, freshly resected PDX tumors were cut into small pieces, and incubated for 30 min at 37 °C in papain solution (10 units/mL, Worthington, LS003126) containing N-acetyl-L-cysteine (160 μg/ml, Sigma-Aldrich, A9165) and DNase I (12 μg/ml, Sigma-Aldrich, DN25), dissociated to single cells by gentle pipetting. Red blood cells in the tumor cell suspension were removed by incubating in RBC Lysis buffer (Stem Cell labs # 07850) at 37 °C for 2 min, followed by rinsing in DPBS- BSA. Cells were filtered using a 40 µm strainer, counted and viability assessed to be above 80%. Cells were plated at 1000 cells/well in 384 well plates in 50% Dulbecco’s Modified Eagle Medium/Nutrient Mixture F12 (DMEM/F12) plus 50% Neurobasal-A Medium supplemented with B-27 supplement (without vitamin A) 2 mmol/L glutamine, 1 mmol/L Sodium Pyruvate, 100 μmol/L non-essential amino acids (NEAA), 10 μmol/L HEPES, 20 ng/mL basic fibroblast growth factor, and 20 ng/mL epidermal growth factor. Serially diluted RBC1HI was added at a final concentration of 40–0.006 µM immediately to the plated cells, with DMSO as negative control and incubated for 72 h. Cell viability at the end of incubation was measured using Cell Titer-Glo Luminescent Cell Viability Assay 2.0 (Promega) with PHERAstar FSX microplate reader. Raw values were converted to cell viabilities and data analyzed using Prism 10 to generate dose-response curves and obtain IC50 values^49^.

### Ex vivo degradation assay in medulloblastoma PDX cells

KBTBD4-PR mutant PDX (ICB1572) tumor was freshly isolated from mouse cerebellum, dissociated to a single cell suspension and plated at 1x10^6^ cells/well in a 6 well plate in 50% Dulbecco’s Modified Eagle Medium/Nutrient Mixture F12 (DMEM/F12) plus 50% Neurobasal-A Medium supplemented with B-27 supplement (without vitamin A) 2 mmol/L glutamine 1 mmol/L Sodium Pyruvate, 100 μmol/L non-essential amino acids (NEAA), 10 μmol/L HEPES, 20 ng/mL basic fibroblast growth factor, and 20 ng/mL epidermal growth factor. Cells were immediately dosed with the HDAC1/2 inhibitor RBC1HI or the NEDD8-activating enzyme (NAE) inhibitor MLN4924 and incubated at 95% humidity and 5% CO_2_. Cells were collected 24 h later, lysed in RIPA buffer, and immunoblotting was performed. Immunoblot images were captured using the LICOR Odyssey CLX Imaging system.

### Cryo-EM sample preparation and data collection

To assemble the complex of KBTBD4-UM171-LHC for cryo-EM study, the individually isolated KBTBD4 protein and co-expressed LHC complex were mixed in stoichiometric amounts with 1 mM UM171 added and subsequently applied to the Superose6 increase gel filtration column (Cytiva) in a buffer containing 40 mM HEPES, pH 7.5, 50 mM KCl, 100 mM InsP_6_ (Inositol Hexakisphosphate) and 0.5 mM TCEP (tris(2-carboxyethyl)phosphine). The isolated complex was then crosslinked with 37.5 mM glutaraldehyde at room temperature for 6 min and quenched the reaction with 1 M Tris-HCl pH 8.0. The crosslinked sample was snap-frozen for future use.

To prepare grids for cryo-EM data collection, an QuantiFoil Au R0.6/1 grid (Electron Microscopy Sciences) was glow discharged for 30 sec at 20 mA with a glow discharge cleaning system (PELCO easiGlow). 3.0 μL of the purified KBTBD4-UM171-LHC complex at 0.7 mg/mL was applied to a freshly glow-discharged grid. After incubating in the chamber at 10°C and 100% relative humidity, grids were blotted for 3 sec with a blotting force of zero, then immediately plunge-frozen in liquid ethane using a Vitrobot Mark IV system (Thermo Fisher Scientific). Data collection was carried out on a FEI Titan Krios transmission electron microscope (Thermo Fisher Scientific) operated at 300 kV at the HHMI Janelia Research Campus. Automation scheme was implemented using the SerialEM^50^ software using beam-image shift^51^ at a nominal magnification of 165 K, resulting a physical pixel size of 0.743 Å. The images were acquired on a falcon 4i camera direct detector, with the slit width of Selectris X (Thermo Fisher Scientific) set to be 6 eV. The dose rate was set to 15.39 e^-^/ Å^2^ S, and the total dose of 60 electrons per Å^2^ for each image were fractionated into 60 EER (Electron-event representation) frames. Data were collected in four sessions with a defocus range of 0.8-1.5 μm. In total, 6,839 movies were collected.

### Image processing and 3D reconstruction

In total, 6,839 movies were collected and imported into CryoSPARC^52^ followed by patch motion correction and patch CTF estimation. 4,982 micrographs were kept after filtering the micrographs with CTF parameters and manual inspection. Blob picker job in CryoSPARC was able to pick 1,788,036 particles, which were further extracted and subjected to 2D classification. After five rounds of cleaning by 2D classification, 171,302 particles were selected and subjected to ab-initio reconstruction. Subsequently, all the particles were used for heterogenous refinement. After one extra round of cleaning up by heterogenous refinement, 110,048 particles from good reconstruction were selected to get re-extracted without Fourier cropping. the homogenous refinement and non-uniform refinement^53^, help reach an overall resolution of 3.88 Å. To better optimize map for the KELCH-repeat domain, a soft mask focused on KELCH domain was applied to local refinement, ending up with a further improved resolution to 3.84 Å. More details about the data processing can be found in Extended Data Fig.4.

### Model building and refinement

The initial structural models of the KBTBD4 dimer and the HDAC1-CoREST-ELM-SANT1 complex was predicted with AlphaFold-Multimer in Google ColabFold^54^. Because the overall model did not fit into the 3.84 Å cryo-EM map very well, the structural models of KBTBD4 BTB-BACK domain, KELCH-repeat domain, and HDAC1-CoREST were separately fit into the cryo-EM map using UCSF Chimera^55^. The resulting model was subsequently rebuilt in Coot^56^ based on the protein sequences and the EM density and was further improved by real-space refinement in PHENIX^57,58^. The structure figures were made using PyMOL^59^.

### Data availability

Coordinates and density map of the KBTBD4-PR-LHC-InsP_6_ and KBTBD4-TTYML-LHC-InsP_6_ complexes are deposited with the PDB and the Electron Microscopy Data Bank (EMDB) under the accession codes 8VRT and 8VPQ, and EMD-43487 and EMD-43413, respectively. All other data are available in the manuscript and the source data files.

---

**Extended Data Figure 1.**
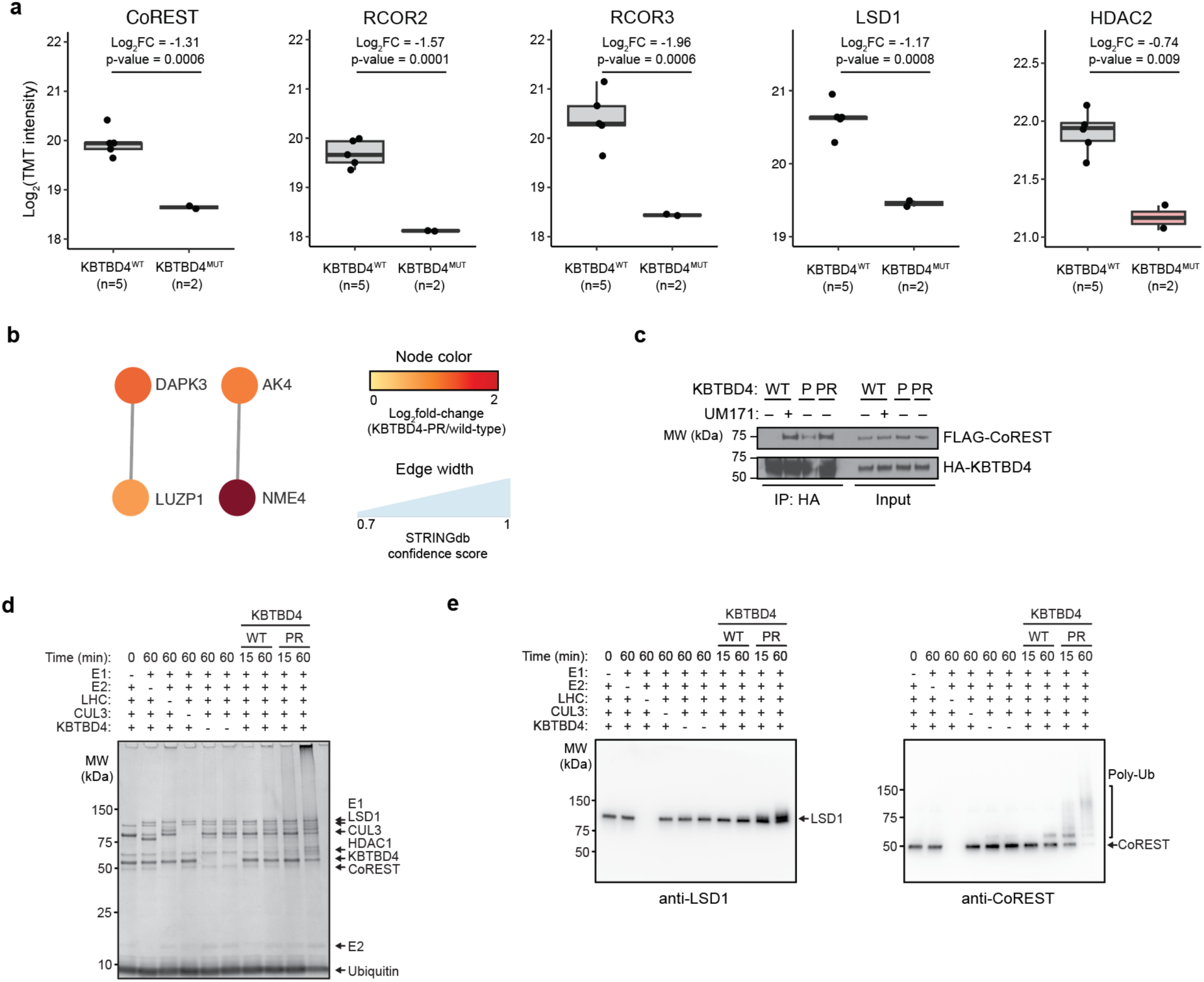
Supporting data for Figure 1. a) Relative protein abundances (log_2_FC(tandem-mass tag (TMT)), *y* axis) in *KBTBD4^WT^*(*n =* 5) and *KBTBD4^MUT^*(*n* = 2) PDX models for selected proteins. P values were obtained using limma’s linear models with empirical Bayes. Box plots show the median and interquartile range, with whiskers extending to 1.5× the interquartile range. b) Protein STRING network showing proteins enriched in *KBTBD4^MUT^* (*n =* 2) versus WT (*n =* 5) PDX models. Node color scale depicts log_2_(fold-change) protein abundance in mutant versus WT models. Edges indicate PPIs, and the width scale depicts confidence of the PPI for the nodes. c) Immunoblots of HA IP in the presence of UM171 (1 µM, 1 h) or DMSO, and MLN4924 (1 µM, 3 h) from 293T cells transfected with FLAG-CoREST and indicated HA-KBTBD4 variant constructs. d) Coomassie staining for in vitro ubiquitination assays of either CRL3^KBTBD4-WT^ and CRL3^KBTBD4-PR^ with LHC. e) Immunoblots for LSD1 (left) and CoREST (right) of in vitro ubiquitination assays of either CRL3^KBTBD4^ and CRL3^KBTBD4-PR^ with LHC. Results in **c** are representative of two independent experiments. Results of **d** and **e** are representative of three independent experiments.

**Extended Data Figure 2.**
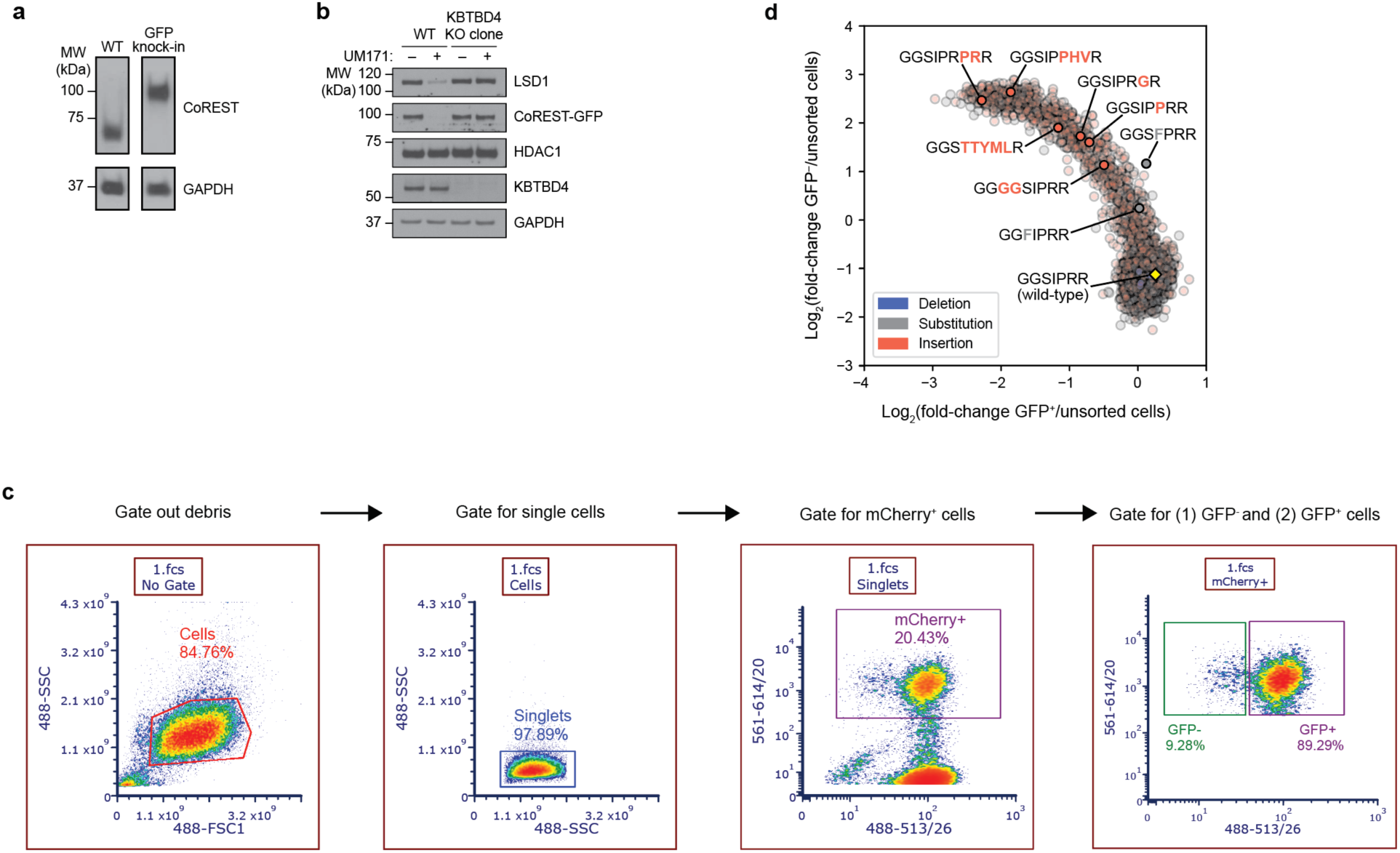
Flow cytometry analysis for CoREST-GFP degradation. a) Immunoblot showing CoREST and GAPDH for WT and CoREST-GFP K562 cells. b) Immunoblot showing CoREST-GFP, HDAC1, KBTBD4, and GAPDH in WT K562 CoREST- GFP knock-in cells and a clonal cell line with KBTBD4 knockout in the presence or absence of UM171 (1 µM, 6 h). c) Representative gating scheme for flow cytometric analysis of CoREST-GFP degradation by KBTBD4. GFP fluorescence and mCherry fluorescence were monitored on the FITC and PE- Texas Red channels, respectively. Data shown is from the deep mutational scanning. d) Scatterplot showing fold-change enrichment of KBTBD4 variants in GFP^+^ (*x-*axis) and in GFP^−^ cells (*y-*axis) versus unsorted cells, normalized to WT. Deletion, substitution, and insertion variants are colored in blue, gray, and red, respectively, and WT is marked by a yellow diamond. MB mutant sequences are labeled. Pearson correlation coefficient *r* = -0.91, p-value = 0.0. Results in **a** and **b** are representative of two independent experiments.

**Extended Data Figure 3.**
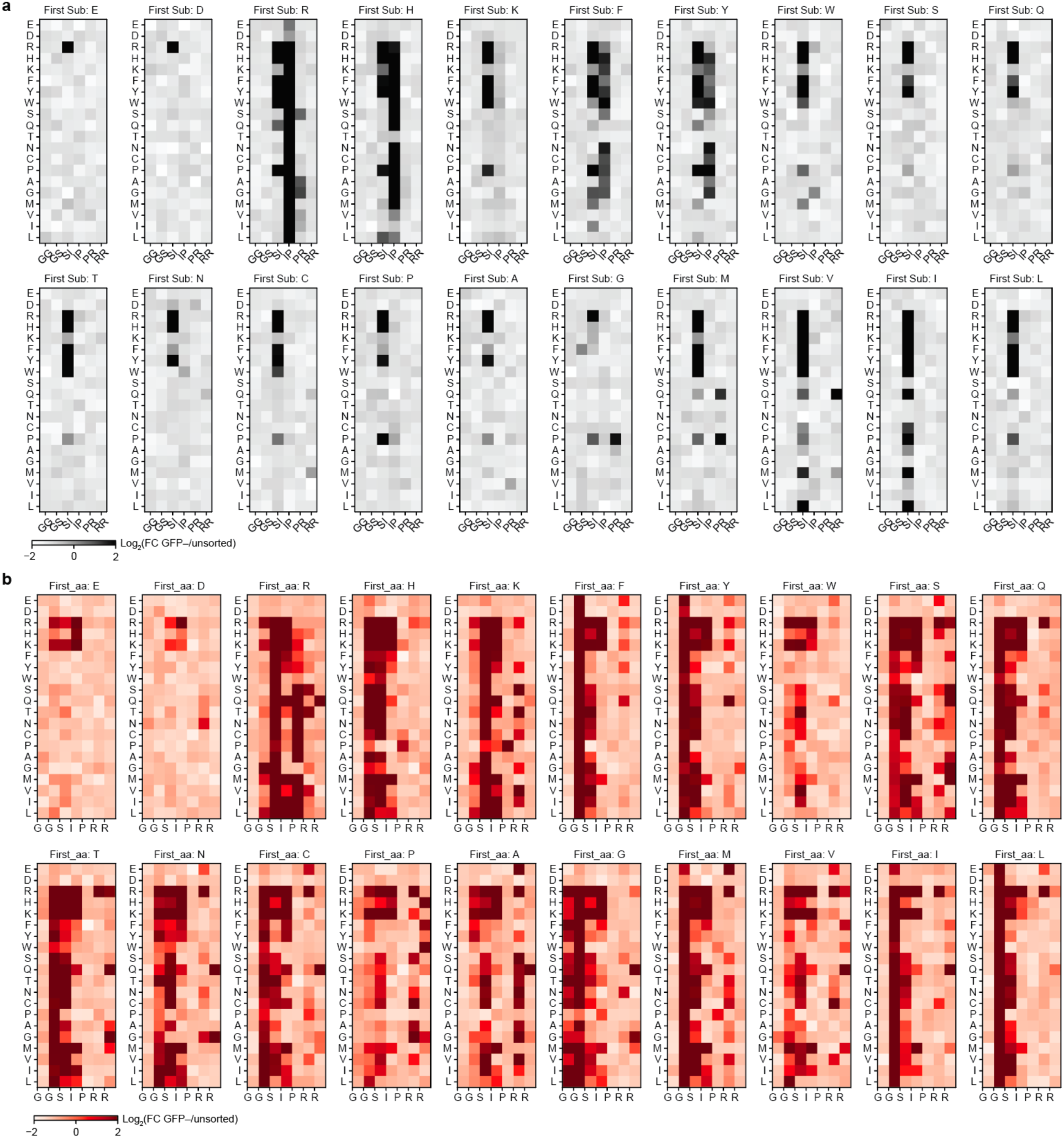
Supporting data for Figure 2. a) Double-substitution deep mutational scanning displayed as heatmaps of fold-change enrichment in GFP^−^ cells for each possible pair of mutated amino acids. b) Double-insertion deep mutational scanning displayed as heatmaps of fold-change enrichment in GFP^−^ cells for each possible pair of mutated amino acids. Color intensity represents mean of *n =* 3 replicates and the overall experiment was performed once.

**Extended Data Figure 4.**
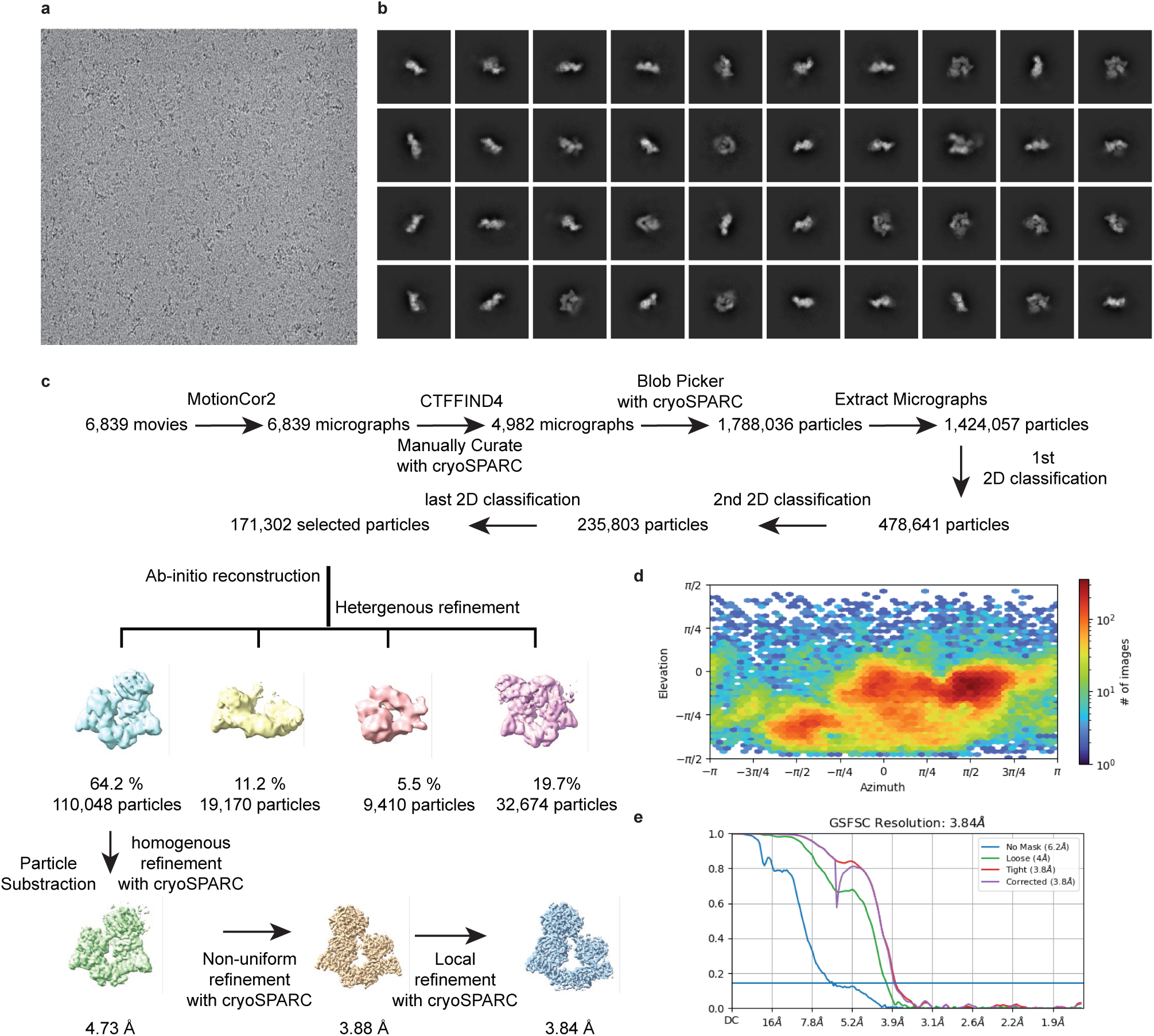
Cryo-EM data processing for the KBTBD4-PR-LHC complex. a) A representative cryo-EM micrograph. b) Typical 2D averages of the cryo-EM dataset; scale bar 10 nm. c) Flowchart of single particle analysis of the KBTBD4-PR-LHC complex. d) Angular distribution of particles used in the final reconstruction. e) Fourier shell correlation (FSC) curves for KBTBD4-PR-LHC. At the Gold-standard threshold of 0.143, the resolution is 3.89 Å.

**Extended Data Figure 5.**
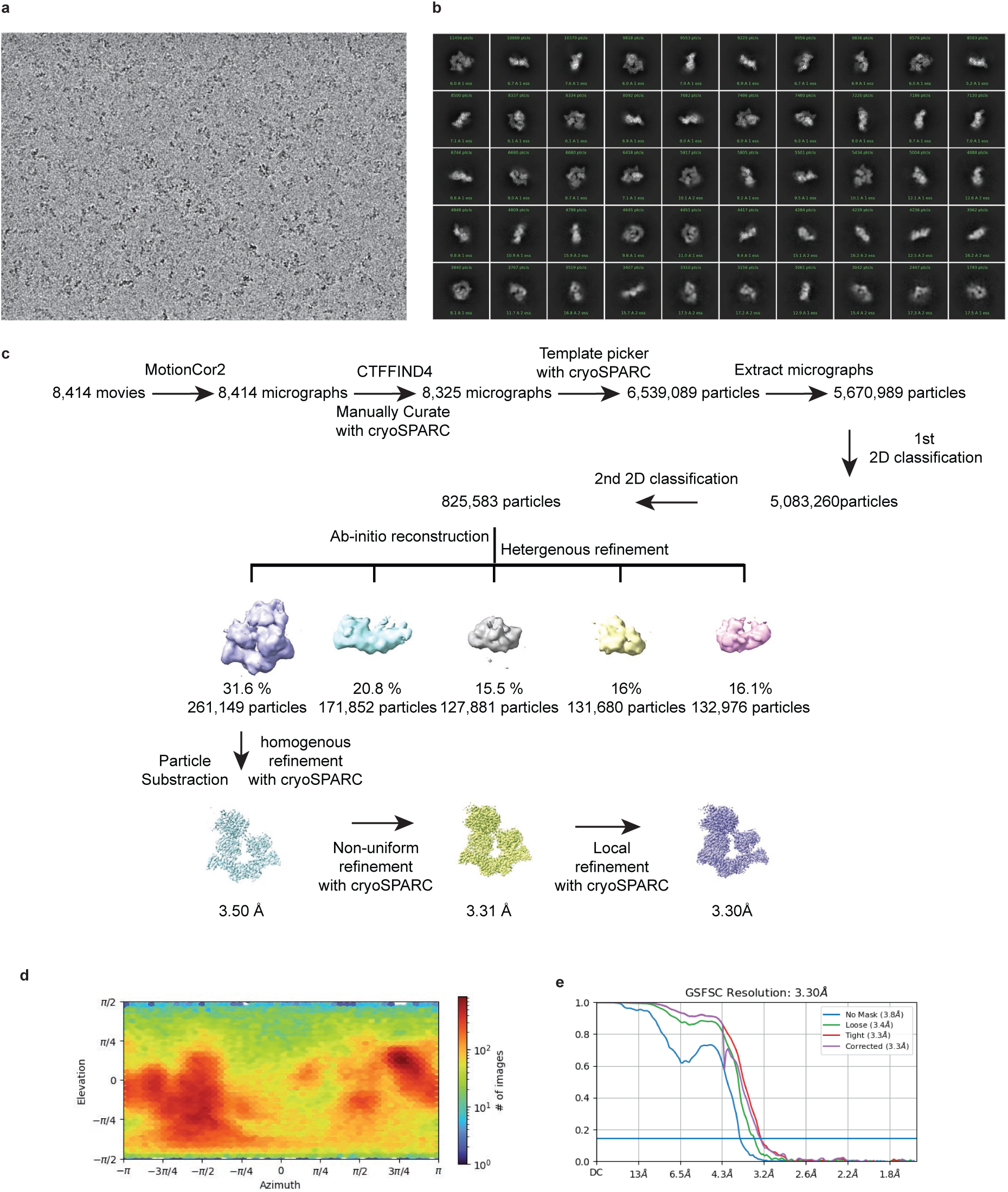
Cryo-EM data processing for the KBTBD4-TTYML-LHC complex. a) A representative cryo-EM micrograph. b) Typical 2D averages of the cryo-EM dataset; scale bar 10 nm. c) Flowchart of single particle analysis of the KBTBD4-TTYML-LHC complex. d) Angular distribution of particles used in the final reconstruction. e) Fourier shell correlation (FSC) curves for KBTBD4-TTYML-LHC. At the Gold-standard threshold of 0.143, the resolution is 3.30 Å.

**Extended Data Figure 6.**
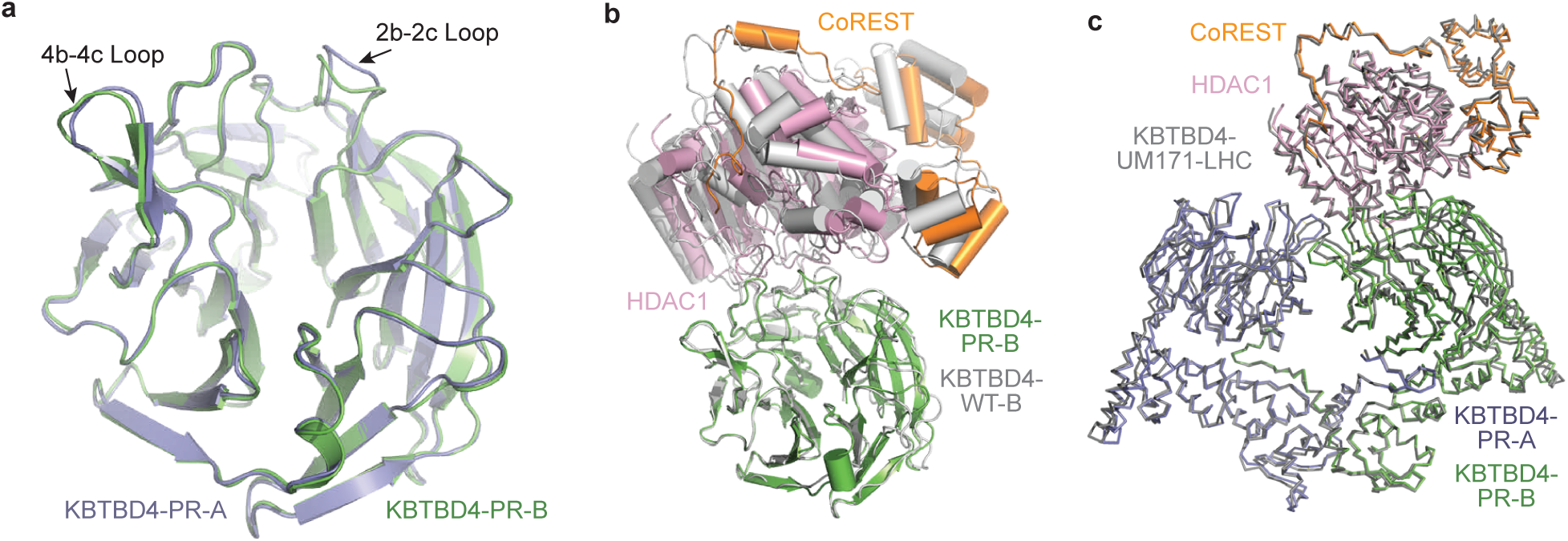
Structural analysis of KBTBD4-PR-LHC complex. a) Superposition of the KELCH-repeat domains of the two KBTBD4-PR protomers (KBTBD4- PR-A: slate; KBTBD4-PR-B: green) in complex with HDAC1. Noticeable structural differences at 2b-2c and 4b-4c loops are indicated. b) A comparison of the relative positions between KBTBD4 β-propeller and the HDAC1-CoREST complex in the KBTBD4-UM171-LHC and KBTBD4-PR-LHC complex structures. The two complex structures are superimposed via the KELCH-repeat domain of KBTBD4. All subunits in the KBTBD4-UM171-LHC complex are colored in gray. UM171 and InsP_6_ are not shown. c) Superposition of the overall complex structures of KBTBD4-UM171-LHC and KBTBD4-PR- LHC. All subunits in KBTBD4-UM171-LHC are colored in gray.

**Extended Data Figure 7.**
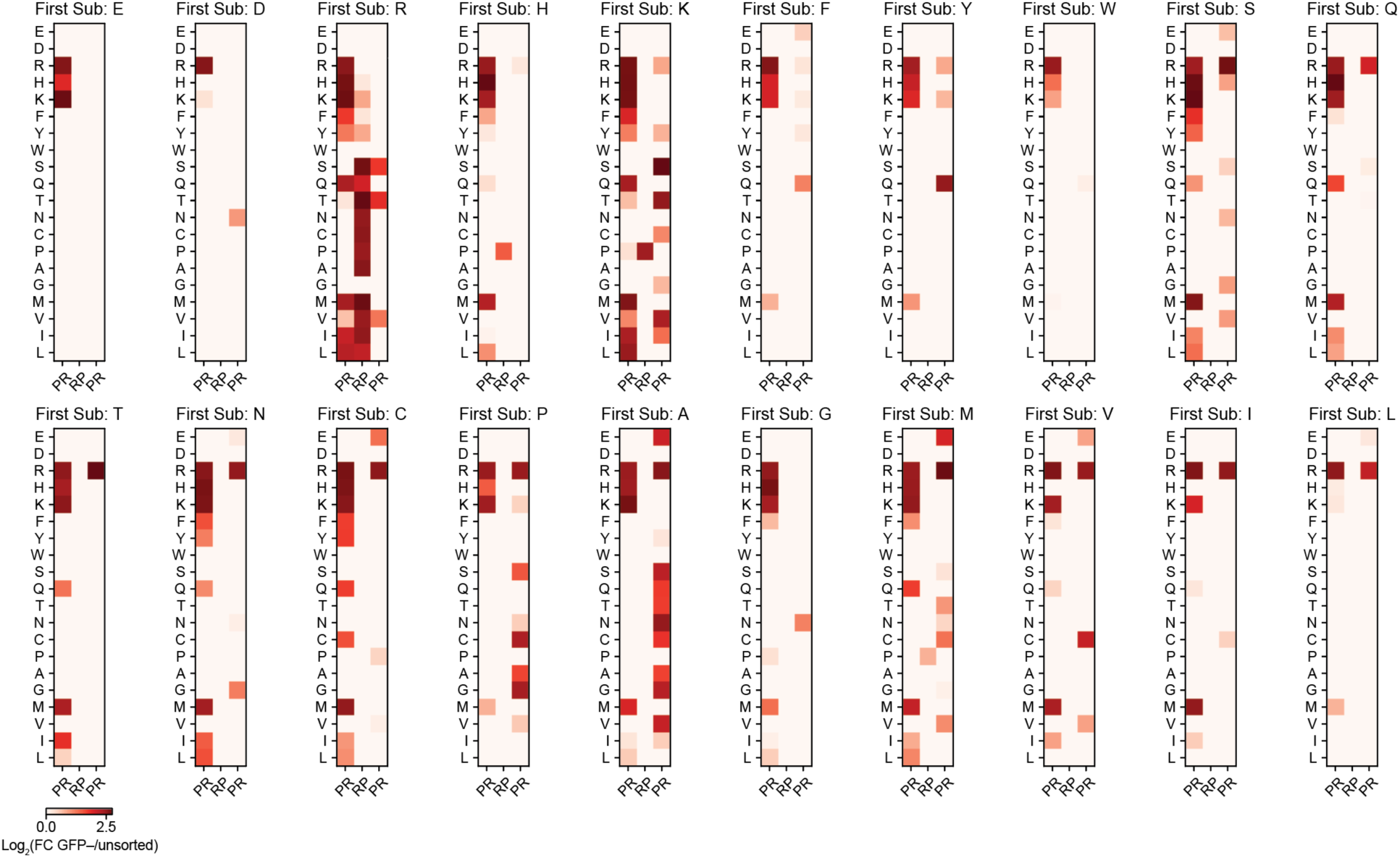
Pairwise saturation mutagenesis of the PRPR sequence of KBTBD4-PR. Double-substitution deep mutational scanning displayed as heatmaps of fold-change enrichment in GFP^−^ cells for each possible pair of mutated amino acids in the KBTBD4-PR PRPR sequence. Color intensity represents mean of *n =* 3 replicates and the overall experiment was performed once.

**Extended Data Table 1.**
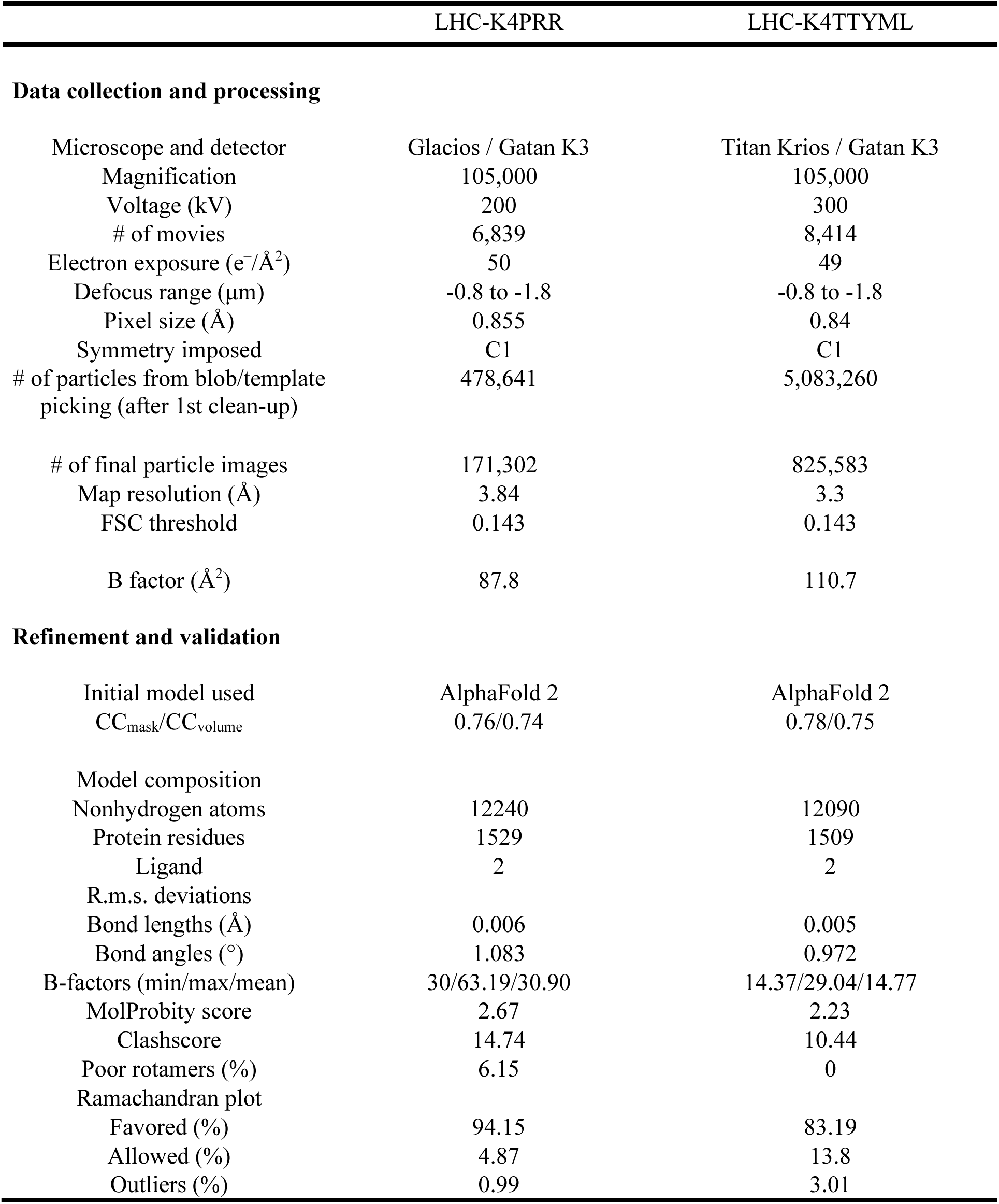
Cryo-EM data collection, refinement, and validation statistics.

|  | LHC-K4PRR | LHC-K4TTYML |
| --- | --- | --- |
| **Data collection and processing** |  |  |
| Microscope and detector | Glacios / Gatan K3 | Titan Krios / Gatan K3 |
| Magnification | 105,000 | 105,000 |
| Voltage (kV) | 200 | 300 |
| # of movies | 6,839 | 8,414 |
| Electron exposure ($e^-/\text{\AA}^2$) | 50 | 49 |
| Defocus range ($\mu\text{m}$) | -0.8 to -1.8 | -0.8 to -1.8 |
| Pixel size ($\text{\AA}$) | 0.855 | 0.84 |
| Symmetry imposed | C1 | C1 |
| # of particles from blob/template picking (after 1st clean-up) | 478,641 | 5,083,260 |
| # of final particle images | 171,302 | 825,583 |
| Map resolution ($\text{\AA}$) | 3.84 | 3.3 |
| FSC threshold | 0.143 | 0.143 |
| B factor ($\text{\AA}^2$) | 87.8 | 110.7 |
| **Refinement and validation** |  |  |
| Initial model used | AlphaFold 2 | AlphaFold 2 |
| CC<sub>mask</sub>/CC<sub>volume</sub> | 0.76/0.74 | 0.78/0.75 |
| Model composition |  |  |
| Nonhydrogen atoms | 12240 | 12090 |
| Protein residues | 1529 | 1509 |
| Ligand | 2 | 2 |
| R.m.s. deviations |  |  |
| Bond lengths ($\text{\AA}$) | 0.006 | 0.005 |
| Bond angles ($^\circ$) | 1.083 | 0.972 |
| B-factors (min/max/mean) | 30/63.19/30.90 | 14.37/29.04/14.77 |
| MolProbity score | 2.67 | 2.23 |
| Clashscore | 14.74 | 10.44 |
| Poor rotamers (%) | 6.15 | 0 |
| Ramachandran plot |  |  |
| Favored (%) | 94.15 | 83.19 |
| Allowed (%) | 4.87 | 13.8 |
| Outliers (%) | 0.99 | 3.01 |

