## Supplementary Notes for "KBTBD4 Cancer Hotspot Mutations Drive Neomorphic Degradation of HDAC1/2 Corepressor Complexes"

**Supplementary Note 1 | Synthetic procedures**

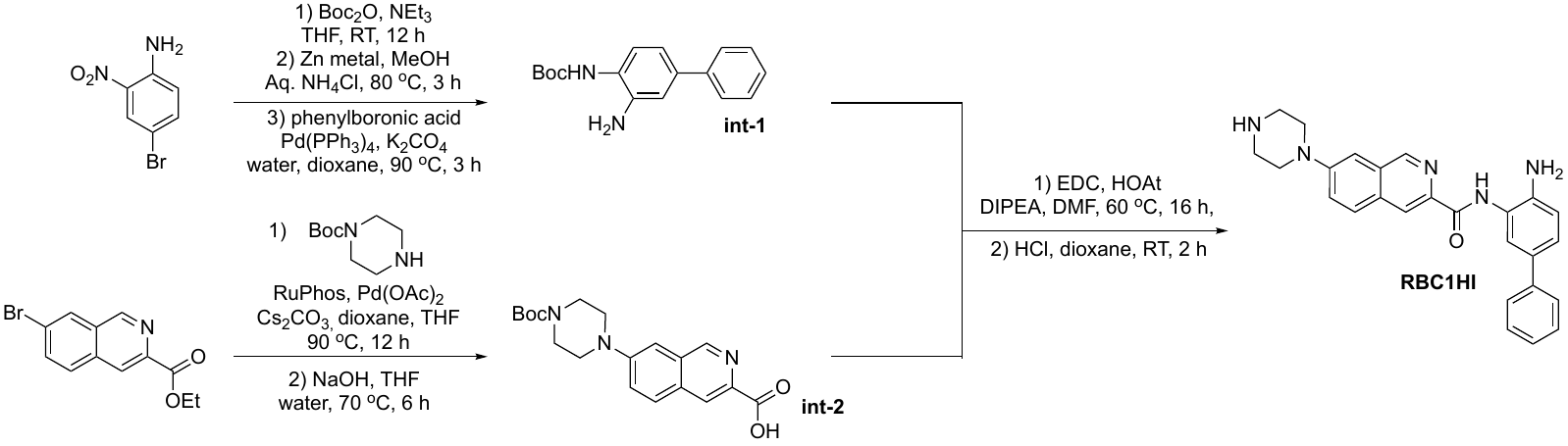

***RBC1HI***

*Synthesis of intermediate 1:*

4-Bromo-2-nitroaniline (5.00 g, 10.9 mmol) was dissolved in THF (100 mL) along with triethylamine (3.03 mL, 21.7 mmol). Subsequently, Boc_2_O (4.74 g, 21.7 mmol) was added portion wise. The resulting solution was stirred at room temperature for 12 hours, followed by concentration. The concentrated solution was redissolved in ethyl acetate (EtOAc) and transferred to a separatory funnel. After washing the crude mixture with water, it was dried over magnesium sulfate, concentrated again, and utilized without additional purification.

tert-butyl (4-bromo-2-nitrophenyl)carbamate (1.00 g, 3.15 mmol) was dissolved in MeOH (50 mL) and saturated aqueous ammonium chloride (15 mL). Zinc powder (1.02 g, 15.8) was then introduced, and the reaction mixture was heated to 80°C for 3 hours. After filtration through celite, the solution was concentrated. The resulting crude mixture was redissolved in ethyl acetate (EtOAc), washed with water, dried over magnesium sulfate, and concentrated once more. Purification by column chromatography on silica gel yielded tert-butyl (2-amino-4-bromophenyl)carbamate (81%, 0.724 g) as a yellow solid.

A mixture of tert-butyl (2-amino-4-bromophenyl)carbamate (1.00 g, 3.48 mmol), phenylboronic acid (0.506 g, 4.18 mmol), and K_2_CO_3_ (2M in water, 10 mL) in dioxane (20 mL) underwent degassing with N_2_, followed by the addition of Pd(PPh_3_)_4_ (0.121 g, 0.105 mmol). The reaction mixture was stirred at 90°C for 6 hours. After cooling, the crude mixture was extracted with ethyl acetate (EtOAc), washed with water and brine. Purification through column chromatography on silica gel resulted in the isolation of tert-butyl (3-amino-[1,1'-biphenyl]-4-yl)carbamate (95%, 0.940 g) as a brown solid.

*Synthesis of intermediate 2:*

The pre-complexation of Pd(OAc)_2_ (0.240 g, 0.300 mmol) and RuPhos (0.600 g, 0.400 mmol) proceeded by stirring in 5 mL of degassed THF at 40 ^o^C for 1h. A mixture comprising ethyl 7-bromoisoquinoline-3-carboxylate (1.00 g, 3.57 mmol), tert-butyl piperazine-1-carboxylate (0.798 g, 4.29 mmol and Cs_2_CO_3_ (1.74 g, 5.36 mmol) in degassed dioxane (30 mL) were added to the activated catalyst flask and stirred at 90 °C under a nitrogen atmosphere overnight. After cooling, the mixture was filtered and concentrated to yield a residue. Subsequent purification through column chromatography produced ethyl 7-(4-(tert-butoxycarbonyl)piperazin-1-yl)isoquinoline-3-carboxylate (94%, 0.646 g) as a yellow solid.

A solution of ethyl 7-(4-(tert-butoxycarbonyl)piperazin-1-yl)isoquinoline-3-carboxylate (0.500 g, 1.30 mmol) in a mixture of MeOH (5 mL) and THF (5 mL) was combined with aqueous 1M NaOH (5 mL) and stirred at 60 °C for 3 hours. The resulting mixture was concentrated to obtain a residue. Water (20 ml) was then added to the residue, and the pH was acidified using 2M HCl. A yellow solid precipitated, which was separated by filtration, washed with water (50 ml), and then dried, yielding 7-(4-(tert-butoxycarbonyl)piperazin-1-yl)isoquinoline-3-carboxylic acid (80%, 0.370 g) as a yellow solid.

*Synthesis of compound RBC1HI:*

A mixture comprising 7-(4-(tert-butoxycarbonyl)piperazin-1-yl)isoquinoline-3-carboxylic acid (0.100 g, 0.280 mmol), tert-butyl (3-amino-[1,1'-biphenyl]-4-yl)carbamate (0.110 g, 0.386 mmol), EDC (0.0738 g, 0.386 mmol), HOAt (0.0524 g, 0.386 mmol) and DIPEA (0.122 mL, 0.702 mmol) in DMF (2 mL) was stirred at room temperature for 16 hours. After evaporation to dryness, the residue underwent purification through silica gel column chromatography, yielding tert-butyl 4-(3-((4-((tert-butoxycarbonyl)amino)-[1,1'-biphenyl]-3-yl)carbamoyl)isoquinolin-7-yl)piperazine-1-carboxylate (80%, 0.139 g) as a yellow solid.

tert-butyl 4-(3-((4-((tert-butoxycarbonyl)amino)-[1,1'-biphenyl]-3-yl)carbamoyl)isoquinolin-7-yl)piperazine-1-carboxylate (was treated with 4N HCl in dioxane (5 mL) and stirred at room temperature for 2 hours, then concentrated to dryness. The resulting solid was washed with ether (20 ml) and subjected to a C18 column chromatography to provide RBC1HI HCl salt (64%, 0.070 g) as a yellow solid. ^1^H NMR (400 MHz, METHANOL-D4): δ 9.45 (s, 1H), 8.96 (s, 1H), 8.24 (d, J = 9.3 Hz, 1H), 8.01 (d, J = 8.5 Hz, 1H), 7.89 (d, J = 2.1 Hz, 1H), 7.78 (m, 2H), 7.71 (m, 2H), 7.63 (d, J = 8.3 Hz, 1H), 7.50 (m, 2H), 7.43 (m, 1H), 3.80 (m, 4H), 3.48 (m, 4H). ^13^C NMR (101 MHz, METHANOL-D4) δ 160.74, 152.03, 146.78, 142.70, 138.40, 131.38, 131.18, 130.96, 130.52, 130.04, 128.87, 128.17, 127.46, 126.73, 126.20, 125.09, 124.86, 124.76, 124.36, 110.11, 44.50, 43.03. HRMS-ESI [M+H]: expected mass: 424.2137 and observed: 424.2133

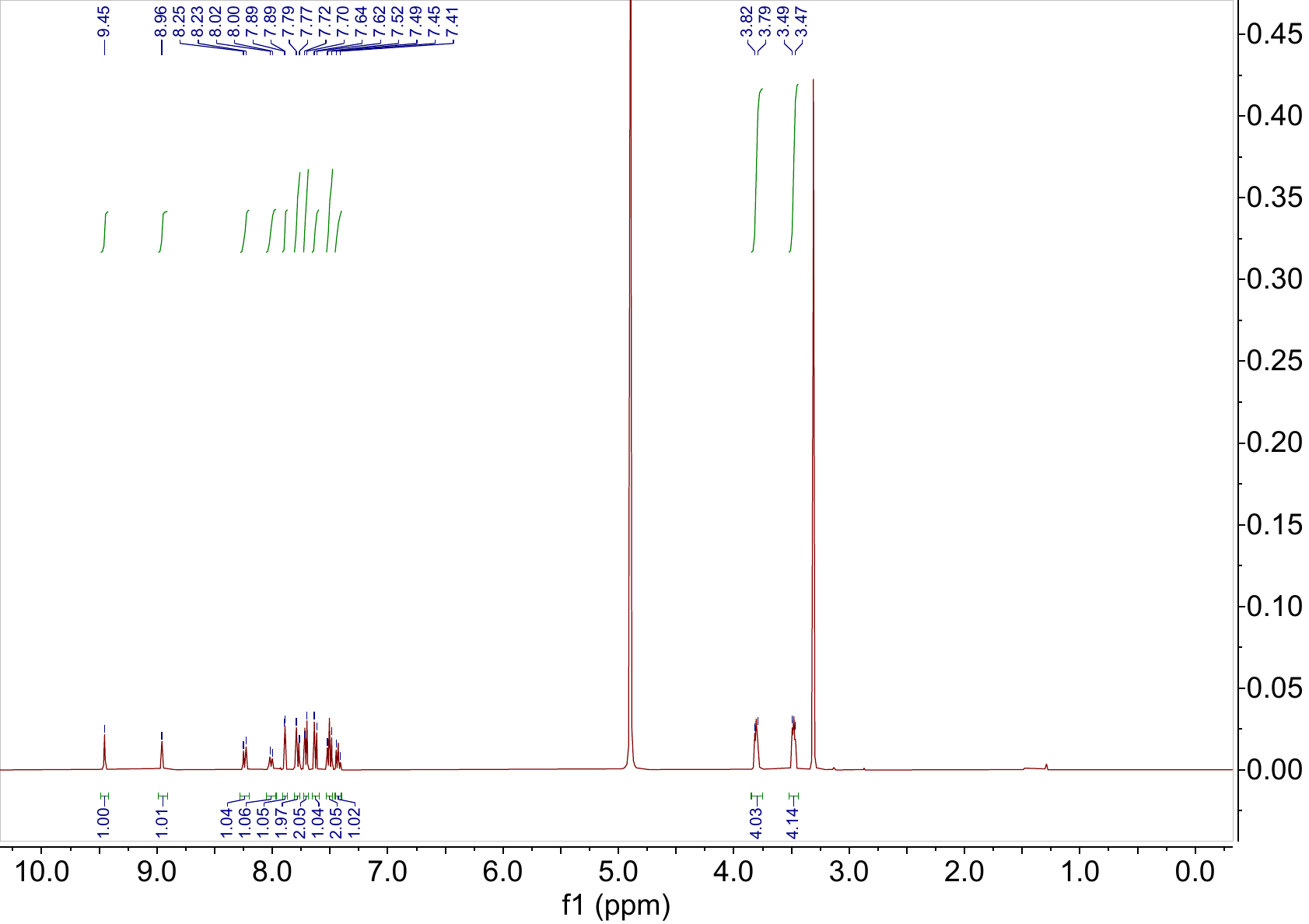

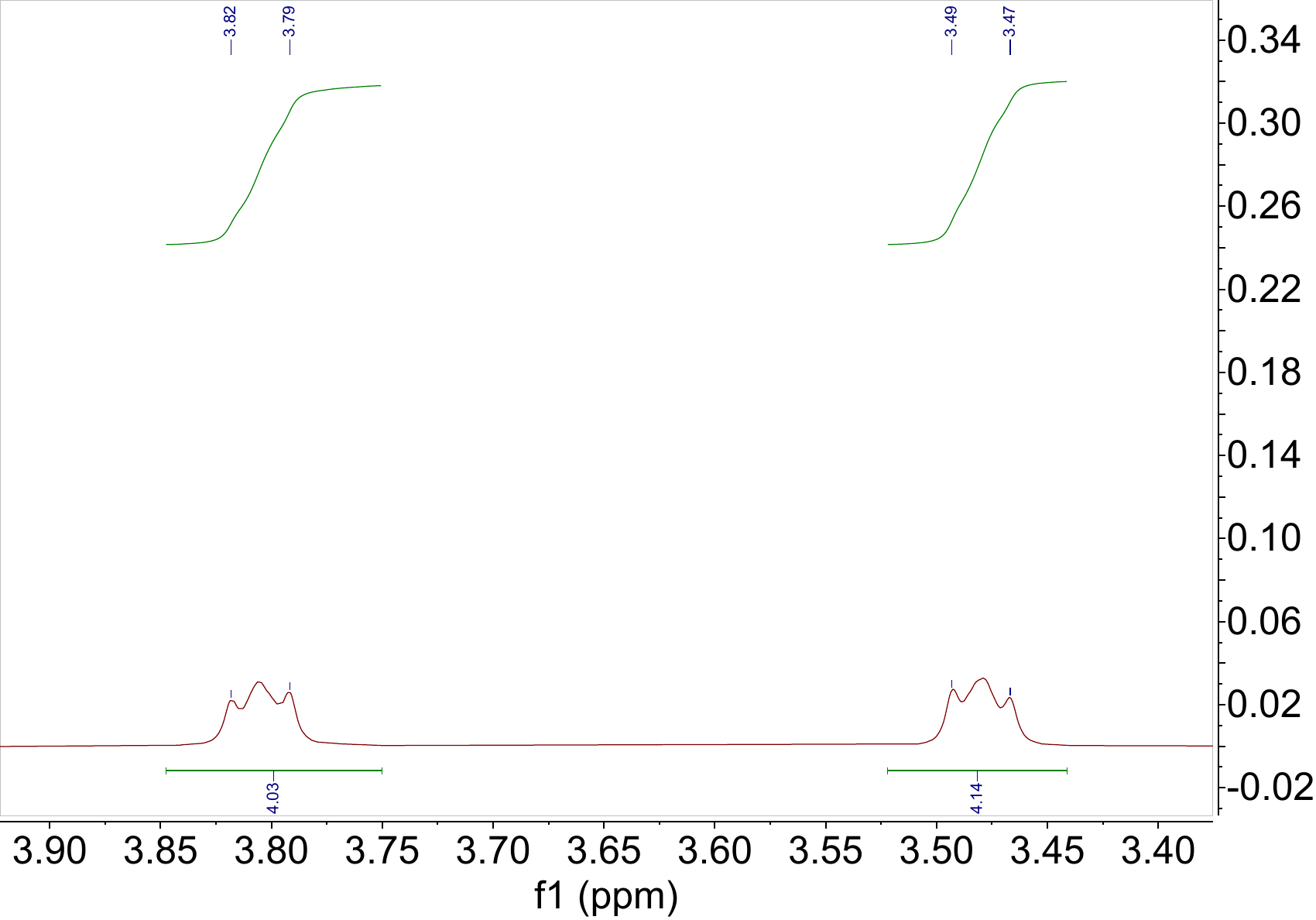

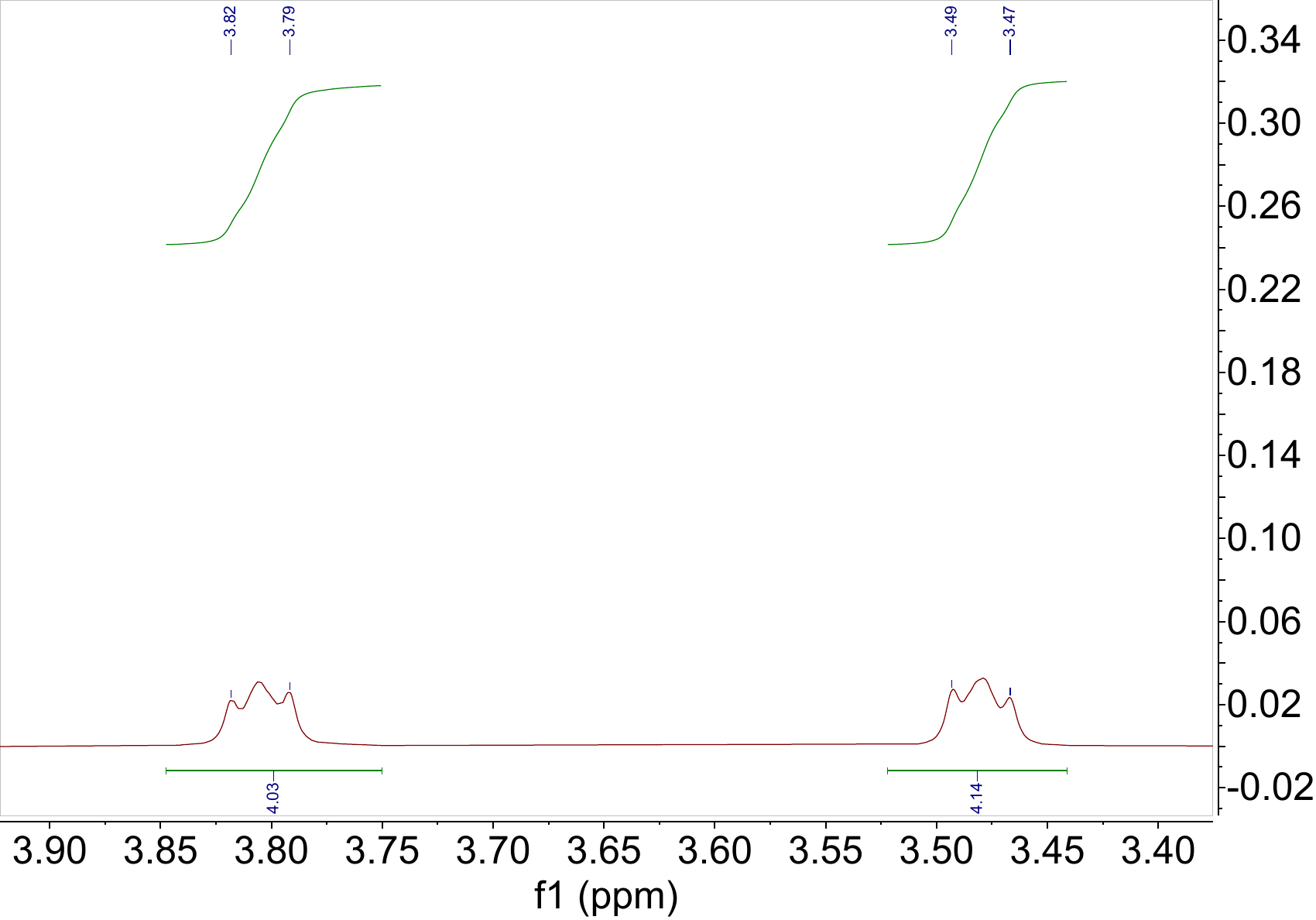

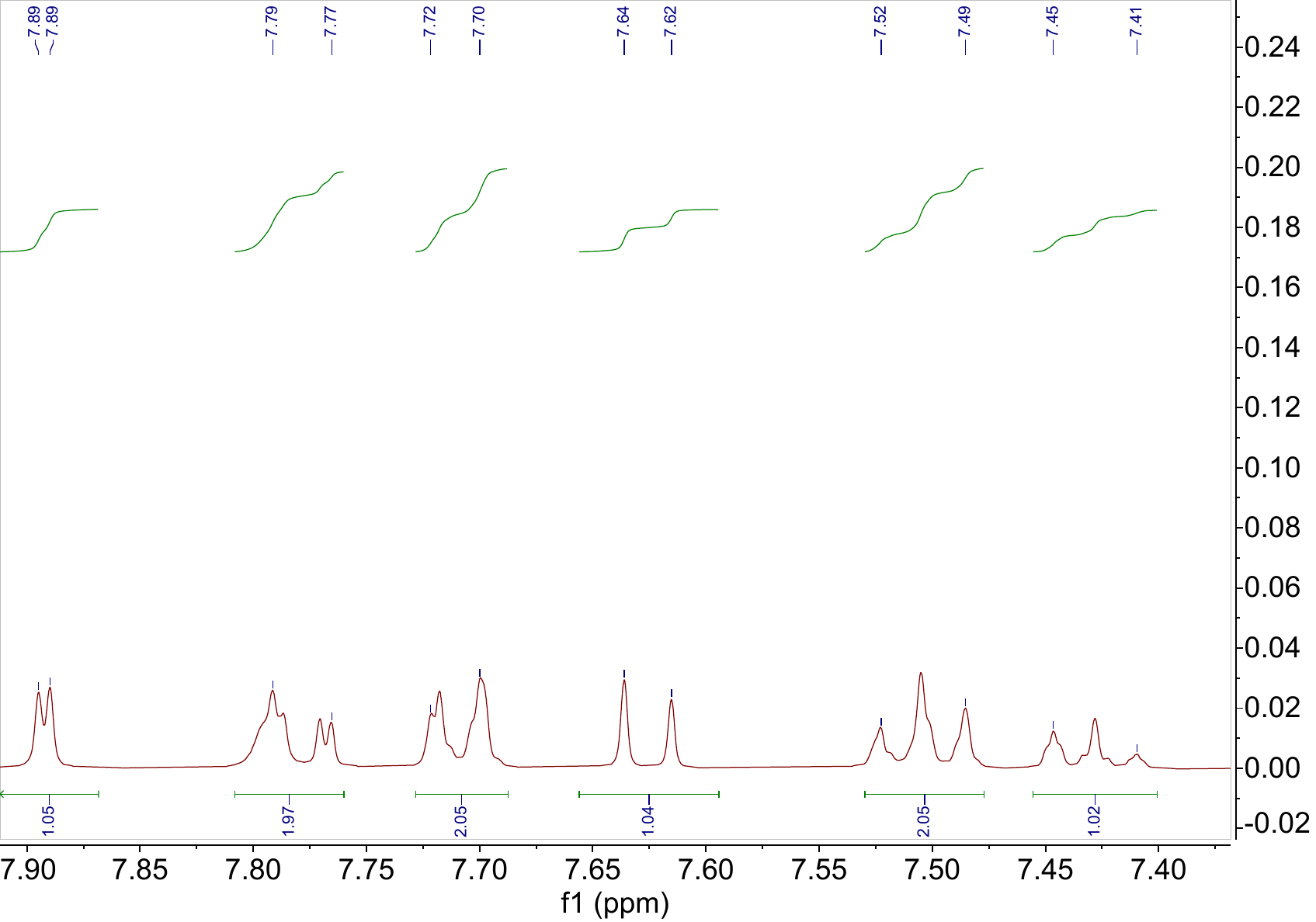

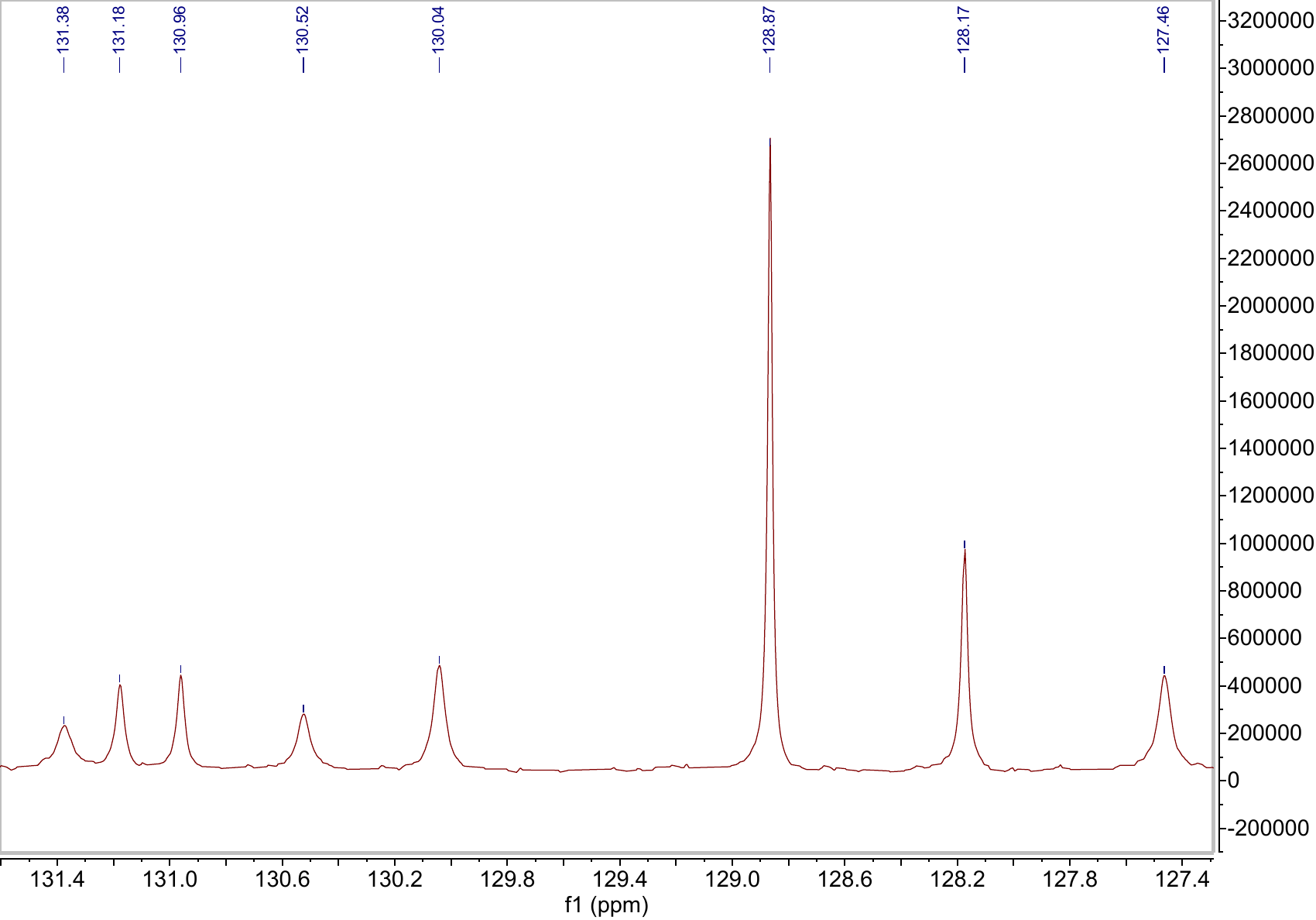

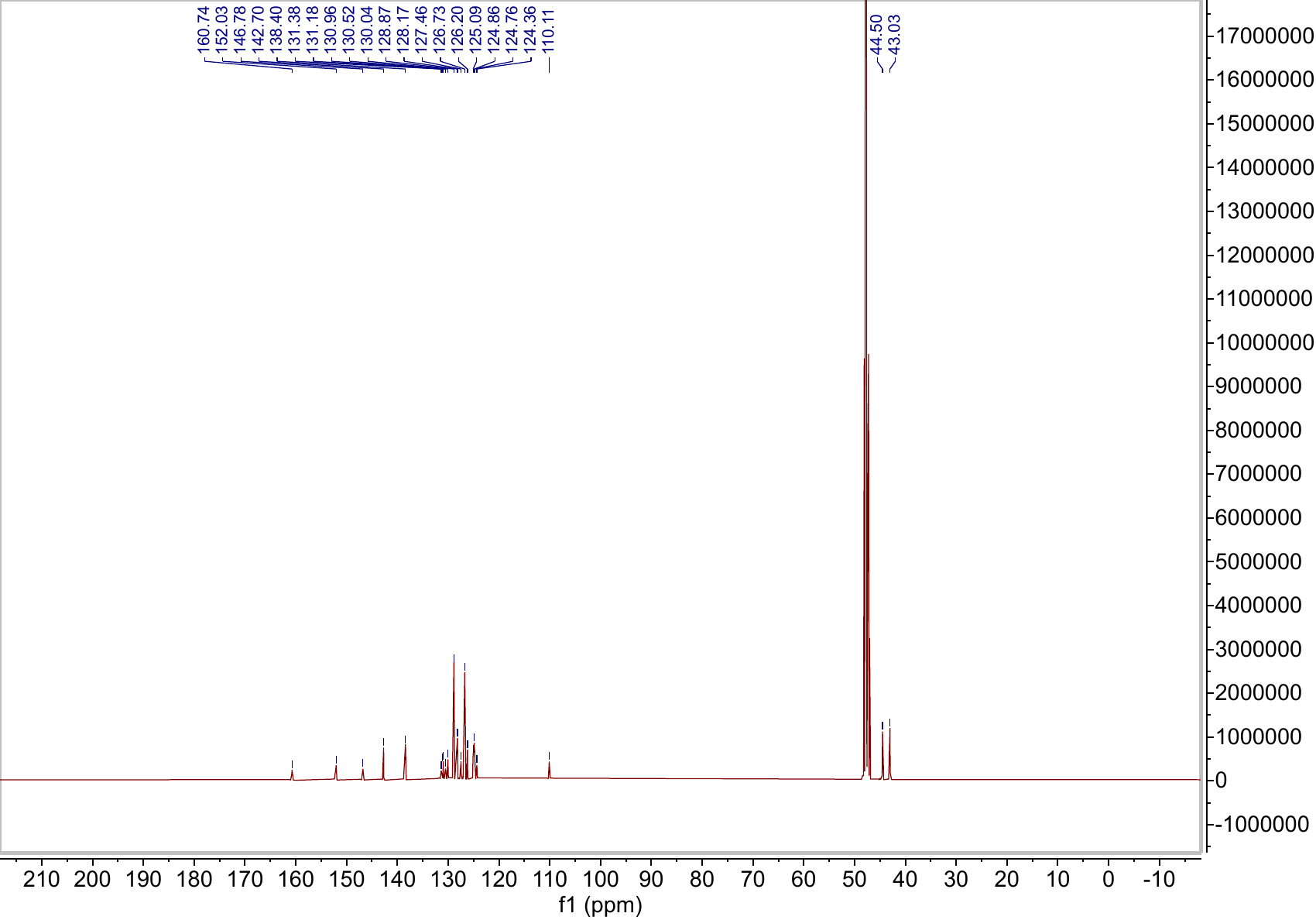
